# Fast Single-Cell MALDI Imaging of Low-Mass Metabolites Reveals Cellular Activation Markers

**DOI:** 10.1101/2024.10.15.618420

**Authors:** James L. Cairns, Johanna Huber, Andrea Lewen, Jessica Jung, Stefan J. Maurer, Tobias Bausbacher, Stefan Schmidt, Pavel A. Levkin, Daniel Sevin, Kerstin Göpfrich, Philipp Koch, Oliver Kann, Carsten Hopf

## Abstract

Single-cell MALDI mass spectrometry imaging (MSI) of lipids and metabolites >200 Da has recently come to the forefront of biomedical research and chemical biology, but fast metabolome-preserving methods without paraformaldehyde fixation for analysis of low mass, hydrophilic metabolites (<200 Da) in large cell populations are lacking. Introducing giant unilamellar vesicles (GUVs) as MSI ground truth for cell-sized objects and Monte Carlo reference-based consensus clustering for data-dependent identification of cell subpopulations. The PRISM-MS (**PR**escan **I**maging for **S**mall **M**olecule - **M**ass **S**pectrometry) dual-scan MSI workflow is presented, enabling space-efficient and therefore faster lipid analysis in single GUVs and cells. Beyond lipids, PRISM-MS enables MSI and on-cell MS2-based identification of low-mass metabolites like amino acids or Krebs cycle intermediates involved in stimulus-dependent cell activation. The utility of PRISM-MS is demonstrated through the characterization of complex metabolome changes in lipopolysaccharide (LPS)-stimulated microglial cells and human-induced pluripotent stem cell-derived microglia. Translation of single cell results to endogenous microglia in organotypic hippocampal slice cultures indicates that LPS-activation involves changes of the itaconate-to-taurine ratio and alterations in neuron-to-glia glutamine-glutamate shuttling. The data suggests that PRISM-MS could serve as a standard method in single cell metabolomics, given its capability to characterize larger cell populations and low-mass metabolites.

## 1. Introduction

Recent advances in spatial- and mass resolution, state-of-the-art machine learning and in on-tissue tandem-MS (MS2) fragmentation analysis for metabolite identification and increased molecular specificity in matrix-assisted laser desorption/ionization (MALDI) trapped ion mobility spectrometry (tims) or Fourier transform mass spectrometry imaging (MSI) have propelled MALDI imaging to the forefront of biomedical research^1–4^. MSI has enabled detailed single-cell analysis within populations of cultured cells or mixtures of cell types^5–11^. Furthermore, population statistics of cultured cells has been assessed by MSI at the single-cell level^8–10,12–14^.

These single-cell platforms either scan entire slides by MSI, which is time-consuming, or smaller fields per slide, thus limiting throughput, or microarrays that hold single cells per spot^9,13,15,16^. Alternatively, they use image-guidance, *i.e.*, orthogonal imaging technologies such as fluorescence microscopy or bright field scanning, to identify cell positions first, and then perform directed cell MSI in the second step^8,12,14^. However, current single cell metabolomics platforms focus on lipidomics and only occasionally on metabolites >200 Da^9^, and they do not cover hydrophilic metabolites with masses below 200 Da such as TCA cycle metabolites or amino acid catabolites. Moreover, metabolites can be extracted from cells during paraformaldehyde (PFA)-fixation and subsequent washing or be altered during the microscopic prescanning step that is usually conducted under ambient conditions. With the exception of a recent protein-focused single cell MSI approach^17^, all other platforms use paraformaldehyde (PFA)- fixed cells.

Here, we introduce the PRISM-MS (**PR**escan **I**maging for **S**mall **M**olecule - **M**ass **S**pectrometry) workflow and a software package that omits PFA fixation and combines a fast and metabolite-preserving low spatial resolution (≥100 µm) MSI PreScan with a higher spatial resolution (≤20 µm) MSI DeepScan. Utilizing giant unilamellar vesicles (GUVs) as cell-sized model membrane systems with known molecular composition as an analytical ground truth^18^, we validate Monte Carlo reference-based Consensus Clustering (M3C) as an automatable approach for selection of cell subpopulations. We translate PRISM-MS with M3C into a search for microglial metabolite activation markers in microglial-like cells, hiPSC-derived microglial cell lines and finally in rat organotypic hippocampal slice cultures.

It has been noted for many years that microglia, arguably in conjunction with reactive astrocytes, play a pivotal role in neurodegeneration, in particular in Alzheimer’s disease (AD)^19–22^. Besides surveying and activated microglia, multiple states of microglial cells characterized by distinct, but complex transcriptomic, proteomic and metabolomic signatures coexist^23^. However, most molecular studies to date have focused on transcriptomics and proteomics, whereas microglial cellular metabolomics remains understudied^24–28^. For instance, in-depth transcriptomics analysis of LPS-stimulated cultured primary mouse microglia, *in-vivo* LPS-treated microglia and human induced pluripotent stem cells (hiPSCs) suggested species differences in metabolic reprogramming during induction of inflammatory responses^29^. Striking changes included alterations in cytokine production, glycolysis or the TCA cycle. For instance, conversion of aconitate to α-ketoglutarate was deregulated post-LPS-exposure^29^. During microglial activation aconitate is decarboxylated to itaconate, the immune cell-specific “posterchild of metabolic reprogramming”, to create a pro-inflammatory response^30,31^. Liquid-chromatography (LC-)MS metabolomics, albeit not at a single-cell level, has recently identified metabolite changes associated with microglia activation in the multiple sclerosis-like experimental autoimmune encephalomyelitis (EAE) model in mice^32^. LC-MS metabolomics highlighted changes in amino acid, taurine and itaconate levels – the latter also demonstrated by DESI-MS imaging – and in other metabolites associated with energy metabolism, oxidative stress and mitochondrial function^32^. Taurine is known to reduce microglia activation in brain^33^.

We therefore demonstrate the utility of the PRISM-MS single-cell MALDI imaging platform in discovery and MS2 characterization of cellular markers of LPS-induced microglia activation.

## 2. Results

### 2.1. PRISM-MS for low mass metabolite-preserving and single cell-focused spatial metabolomics

PRISM-MS utilizes two subsequent scans - a PreScan at low spatial resolution (≥100 µm) that can employ any cell marker like FA18:1 (*m/z* 281.25 [M-H]^-^) for fast, whole-slide identification of cell containing spots. A subsequent DeepScan at high spatial resolution (≤20 µm) is restricted to cell-containing PreScan fields (**Figure 1 a, b** and **Figure S1**). For cultures with intermediate cell density (1000 cells/cm²), we noted a 12- fold speed advantage and concomitant reduction of file size (50 instead of 600 GB; **Figure S2 and S3**) compared to whole-slide scans at 20 µm. The more 200 µm PreScan pixels are devoid of cells, the higher the speed and file size advantage. The user-selected pixel-size of the DeepScan is typically ≤20 µm. PRISM-MS works well with both light-transmitting slides (*e.g.,* glass or ITO chamber slides or droplet microarrays^34^) and non-light-transmitting slides (such as gold-coated slides) (**Figure S4 to S6**). Furthermore, PRISM-MS can combine different modalities such as positive and negative ion modes, and it is compatible with on-tissue data-dependent fragmentation analysis for metabolite identification, *e.g.,* MALDI-imaging parallel reaction monitoring-parallel accumulation serial fragmentation (iprm-PASEF)^4^ or MALDI-TIMS-MS2-MSI approaches^2^.

**Figure 1.**
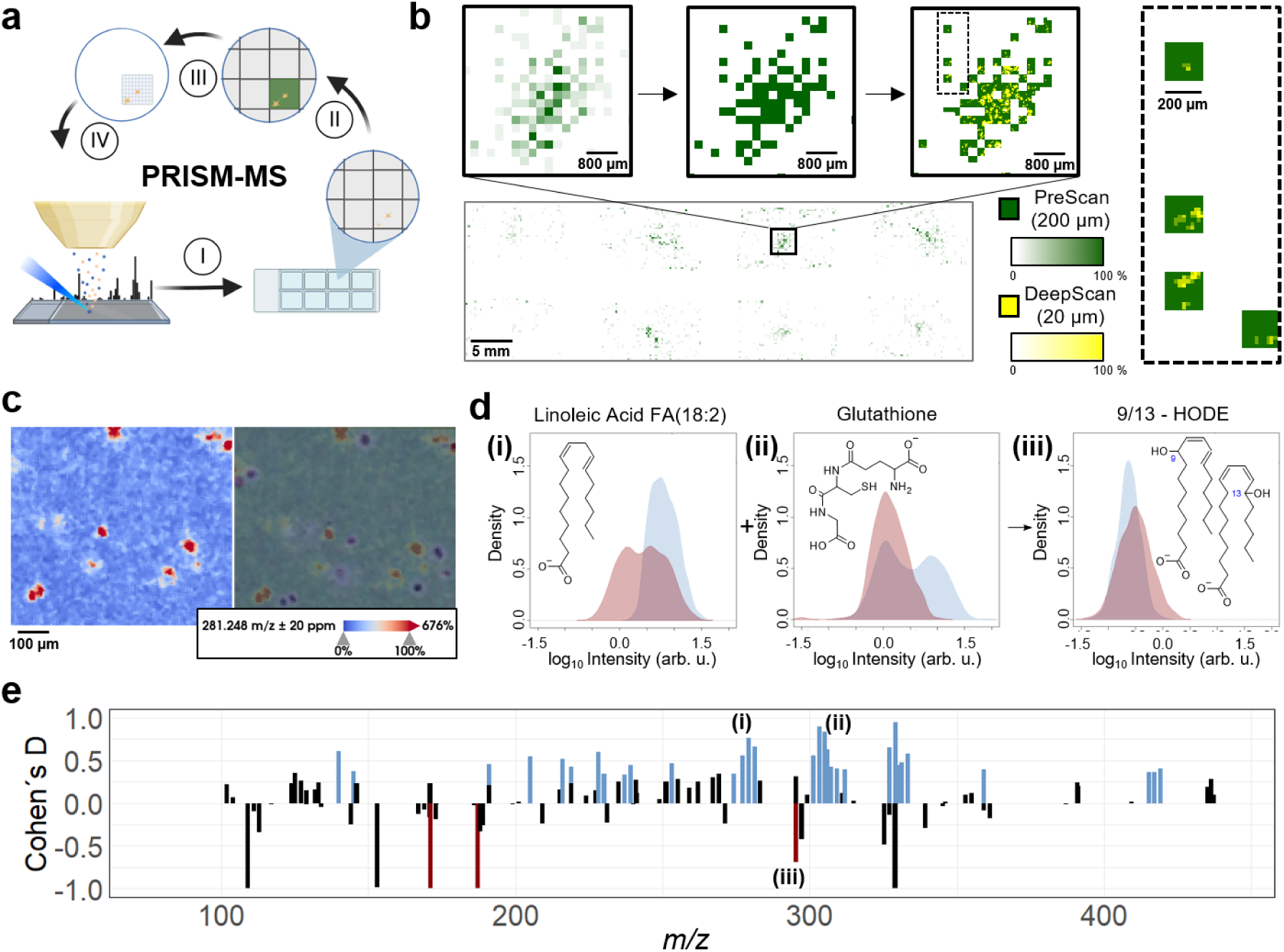
PRISM-MS for fast low mass metabolite-preserving single-cell spatial metabolomics. **a)** Schematic overview of PRISM-MS workflow. (I) Survey PreScan at large (e.g. >100 µm) pixel size; (II) determine cell containing pixels by feature-selective binary image segmentation; (III) definition of cell containing pixels as new measurement regions; (IV) DeepScan at small (e.g. <20 µm) pixel size; followed by data analysis for single cell metabolomics and cluster detection (**Figure S12**). **b)** PRISM-MS example of cultured SIMA9 mouse microglia cells covered with MALDI matrix directly after lyophilization: the PreScan obtains cellular signal intensities per 200-µm pixels (green color scale) (upper panel left), which then get thresholded (upper panel middle) and undergo a subsequent Deep-Scan at 20-µm pixel size to resolve individual cells (upper panel right). Right dashed box: four distinct 200-µm measurement regions with 20 µm DeepScan step size (ion images of *m/z* 281.25 (FA 18:1 [M-H]^-^)). **c)** left panel: 5-µm MSI image of cell marker *m/z* 281.25 (FA 18:1 [M-H]^-^); right panel: overlay of ion image with haematoxylin and eosin (H&E) -stained slide generated after MSI. The registration offset is deliberate, in order to demonstrate cell identification versus H&E ground truth. **d)** Intensity profiles for (i) *m/z* 279.23 (linoleic acid [M-H]^-^), (ii) *m/z* 306.08 (glutathione [M-H]^-^) and (iii) *m/z* 295.23 (9- and 13-hydroxy-octadecadienoic acids (9/13-HODE) [M-H]^-^) obtained by PRISM-MS (blue) versus the optically guided workflow (red). Oxidation of linoleic acid to 9/13-HODE is reduced in PRISM-MS. **e)** PRISM-MS preserves metabolite profiles: average (*N*=3) Comparison of Cohen’s D effect sizes for small molecule *m/z* features obtained by PRISM-MS or by a workflow including 30 min slide exposure to ambient conditions to emulate workflows that capture, e.g., a high resolution optical image before MSI^14^. Metabolites with significantly higher (blue) or lower (red) effect sizes/ intensities in PRISM-MS versus emulated optically guided workflow. Black bars indicate peaks with non-significant (p >0.05; N=3 each for PRISM and emulated optical workflow) differences between the workflows. This figure was partially created using BioRender.

PRISM-MS users can adjust two scanning parameters (**Figure S7**): A) the dilution factor computationally dilates PreScan pixels before the DeepScan, in order to avoid capturing incomplete cells (**Figure S1**). B) The intensity threshold that is commonly defined by the mean intensity for the target *m/z* (*e.g., m/z* 281.25) ± 1 to 2.5 standard deviations. This ensures that most cells seeded on the slide are captured. It is subsequently listed as the number of standard deviations *T_N_*.

Fluorescence scans of Hoechst-stained SIMA9 microglia cells or H&E-stained cells (**Figure 1c**) as a ground truth for absence or presence of cells suggested that false positive/negative rates could effectively be fine-tuned using these parameters (**Figure S7 and S8**), resulting in different file sizes (**Figure S3**). For example, with a dilation factor of 1 and a threshold of *T_N_* =1, the false-positive and false-negative rates for cell ID in MSI were typically 6.5% and 76.7%, respectively. With dilation factor of 1 and threshold *T_N_* =3, the false-positive and false-negative rates for cell ID in MSI were typically 1.9% and 90.1%, respectively.

The driving force behind development of PRISM-MS was the need to cover metabolites with *m/z* below 200 that are of major interest in metabolomics and to improve metabolite preservation throughout sample preparation and imaging (**Figure S9**). Current single cell spatial metabolomics workflows use PFA-fixed cells. This type of fixation may increase imaging quality for single cell lipidomics, but it causes leakage of cytosolic hydrophilic compounds^10^. These metabolites may be reduced by 90% after PFA-fixation^8^. In addition, during optical-guidance-based MSI approaches, extended incubation of cells at room temperature during acquisition of high-resolution microscopic images that define cell positions may also cause undesired metabolome alterations (**Figure 1e**).

In contrast, we chose to fix cultured cells on slides by immediate lyophilization and by covering them with UV-shielding MALDI matrix chemicals and fixing organic solvents as quickly as possible to preserve the metabolomic state. Lyophilization can immobilize molecules inside samples within milliseconds^35^. Comparison of PRISM-MS with a 30 min incubation of unfixed cells at room temperature prior to MSI to emulate an optically guided workflow led to substantially different metabolite profiles, calculated as Cohen’s D, a standardized effect size for assessing the difference between two groups (**Figure 1e**). Cells assessed via microscopic guidance may not represent stable metabolomes, as they are exposed to heat, UV-light (not being shielded by MALDI matrix) and oxygen during extended optical scanning. In PRISM-MS, glutathione depletion, oxidation of linoleic acid to bioactive pathology-associated 9- or 13-hydroxy-octadecadienoic acids (9/13-HODE; **Figure 1d**) and oxidation of additional polyunsaturated fatty acids (**Figure S10**) were all reduced compared to an emulated optically guided workflow^36^. PRISM-MS also led to a slightly higher total number of false-discovery rate (FDR)- controlled metabolite annotations in METASPACE^37^ (**Figure S11**).

As PRISM-MS provides an MSI-only analysis with higher speed and improved metabo- lite preservation compared to microscopy-based imaging, it requires rather low cell culture seeding densities of 1000 cells per cm^2^. The PRISM-MS data processing pipeline i) clusters MSI pixels into “cells”/objects, then ii) distinguishes single cells from cell aggregates/conglomerates and iii) removes the latter from consideration. In such low-density cell cultures, up to seven pixels, most of which contain only a small portion of the cell, can be clustered together to define a single cell, and such cultures typically contain >90% single cells (**Figure S12**).

### 2.2. Giant unilamellar vesicles (GUVs) as analytical ground truth for Monte Carlo reference-based Consensus Clustering (M3C) of cell subpopulations in PRISM- MS

GUVs are cell-sized model membrane systems characterized by their defined lipid constituents. Given that the subset of metabolites present in any set of cells is generally unknown, the lack of that ground truth often complicates the testing of new models or algorithms. We therefore reasoned that GUVs could serve as a (qualitative) ground truth in MSI to validate the PRISM-MS single cell analysis pipeline. We employed two types of single-lipid GUVs composed of 1,2-dioleoyl-sn-glycero-3-phosphocholine (DOPC) and 1,2-dimyristoyl-sn-glycero-3-phosphocholine (DMPC) with heterogeneous size-distributions of up to 50 µm in diameter and a mixture of the two GUV types (**Figure 2a to c** and **Figure S13**)^18^. As a control, we included 1% lissamine rhodamine B-labeled phosphatidylethanolamine in the lipid mixture. Colocalization of fluorescence label and DOPC MSI ion image of dual-labelled GUVs confirmed that the PRISM-MS DeepScan effectively detected and characterized these structures (**Figure 2d**).

**Figure 2.**
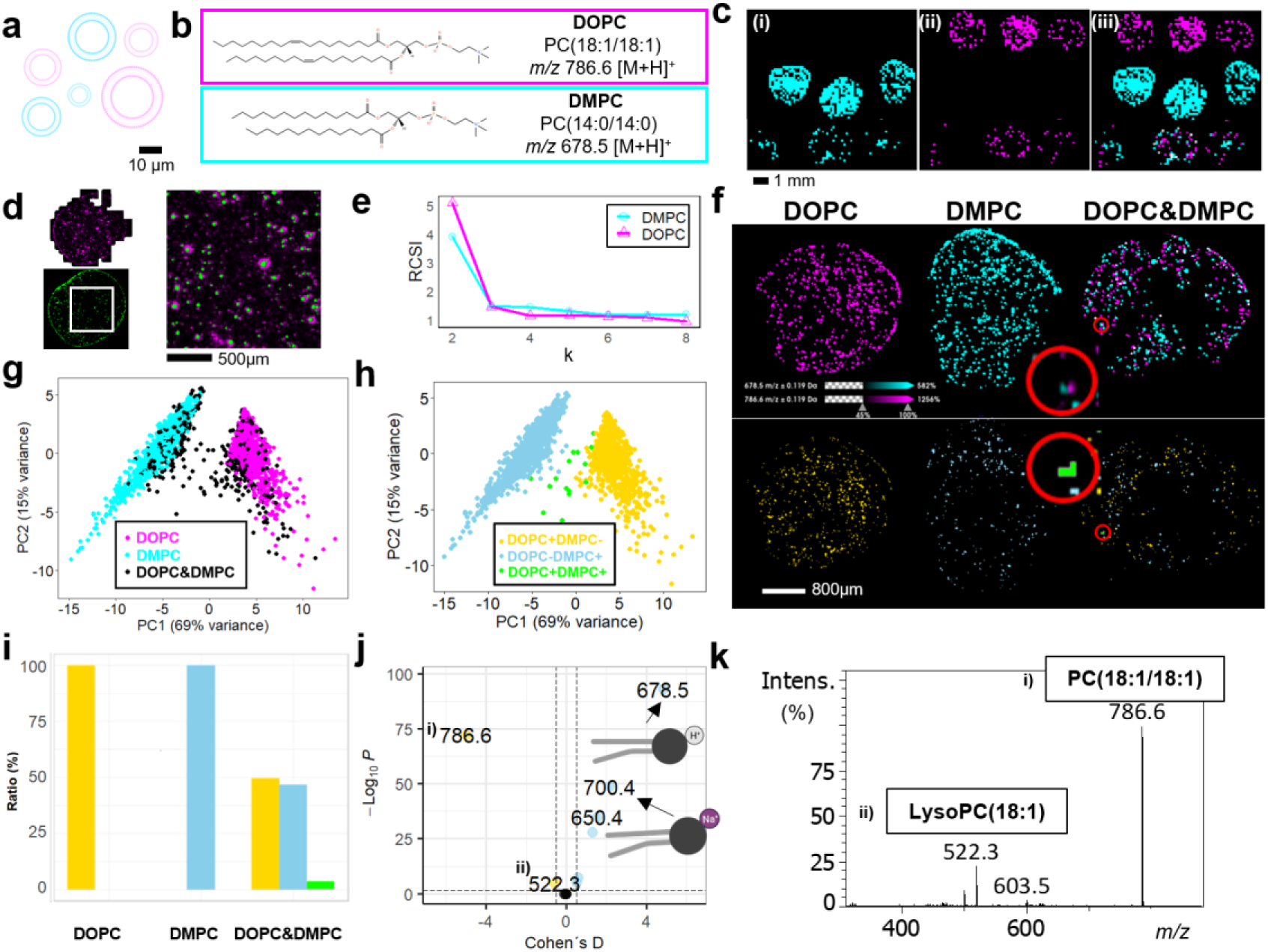
Giant unilamellar vesicles (GUVs) as an analytical ground truth for validation of Monte Carlo Consensus Clustering (M3C). **a)** GUVs are 5-50 µm single-lipid vesicles composed of either DOPC (magenta) or DMPC (cyan). **b)** Chemical structures of protonated DOPC (*m/z* 786.6; phosphatidylcholine PC(18:1/18:1) [M+H]^+^) and DMPC (*m/z* 678.5; PC(14:0/14:0) [M+H]^+^). **c)** Binary thresholded image of the PRISM-MS 200-µm PreScan displayed for (i) DOPC, (ii) DMPC and (iii) a 1:1 mixture of both GUV types. Three spots each were applied onto ITO slides. **d)** Overlay of fluorescence image of lissamine rhodamine B (red; 1% LissRhod-PE included in lipid mixture) label in GUVs, and 20- µm pixel DeepScan of *m/z* 786.6 (DOPC; cyan) **e)** Monte Carlo Consensus Clustering (M3C) for a 1:1 mixture of DOPC and DMPC GUVs on a full ITO chamber slide. Relative Cluster Stability Index (RSCI) suggests k=2, consistent with two classes of GUVs, as the most stable cluster with an overall p< 0.01. **f)** Top row – the ground truth: ion image overlays of *m/z* 786.6/DOPC and *m/z* 678.5.6/DMPC for three conditions: pure DOPC-, pure DMPC- and mixed DOPC/DMPC GUVs. Bottom row – M3C model generation: M3C clustering (k=2) classifies GUVs as DOPC+ (yellow), DMPC+ (blue) or DOPC+/ DMPC+ (gray). **g)** Principal component analysis (PCA) of GUVs; plot of PC1 and PC2 for the three conditions, DOPC (magenta), DMPC (cyan) and mixed DOPC/DMPC (black). **h)** Post-clustering PCA of GUVs categorized by the M3C model enables class assignment for each GUV: DOPC+/DMPC- (yellow), DOPC-DMPC+ (blue), and DOPC+DMPC+ (grey). **i)** M3C model-based population statistics for the three conditions in (g) suggests that 3.5% of GUVs in DOPC/DMPC mixture are classified as DOPC+DMPC+. **j)** Cohen’s D effect sizes versus p-value Volcano Plot for M3C cluster analysis: *m/z* features specific to DMPC-GUV populations, such as *m/z* 678.5 (DMPC [M+H]^+^) and *m/z* 700.5 (DMPC [M+Na]^+^), or specific to DOPC-GUV populations like *m/z* 786.6 (DOPC [M+H]^+^) and the corresponding lysophosphatidylcholine (LPC) *m/z* 522.3. **k)** MS2 spectrum of (i) *m/z* = 786.6 indicating (ii) LPC(18:1) [M+H]^+^ as a likely DOPC fragment in j).

Using the GUV ground truth data, we evaluated a recent clustering algorithm, Monte Carlo reference-based Consensus Clustering (M3C) that had not been used in MSI yet. In popular clustering methods like k-means clustering the number of classes k is either arbitrarily picked or inferred according to the Calinski-Harabasz criterion, Silhouette score or other clustering metrics^38^. M3C was developed to detect heterogeneities in genomic data and to avoid expectation bias^38^. For high dimensional data like genomic or MSI data M3C repeats a statistical test for various k values, for instance, 200 times (as in Monte Carlo simulations) and identifies those k that show significant differences from a homogeneous distribution. The algorithm then evaluates a reference cluster stability index (RCSI) for all significant k values. The most stable k is ultimately used for segmentation^38^.

We analyzed DOPC-only and DMPC-only GUVs and a 1:1 mixture of the two GUV types by PRISM-MS. For both lipid *m/z* tested separately, the RCSI in M3C was highest for k=2 subpopulations suggesting that *m/z* for both DOPC and DMPC were not evenly distributed across the entire population (**Figure 2e** and **f**). This analysis suggested four possible groups and labels: DOPC-/DMPC+ (1), DOPC+/DMPC- (2), DOPC+/DMPC+ (3) and DOPC-/DMPC- (not observed). Even in the mixture, most GUVs were classified as (1) or (2) (**Figure 2g to i**). The small number of vesicles in the mixture classified as positive for both (3) were typically two vesicles in close proximity that were not seperable at 20 µm lateral step size (**Figure 2f, 2i** and **Figure S14**). Most GUVs were represented by a single 20-µm pixel. Even though GUV sizes varied, size differences did not affect their group assignment (**Figure S14**). Annotation of molecular differences between DOPC-/DMPC+ (1) and DOPC+/ DMPC- (2) clusters in METASPACE followed by MS2 fragmentation analysis revealed 13 features including a DOPC fragment, LysoPC (18:1) (**Figure 2j** and **k**). As expected and as a validation of the approach, both lipids including variants such as their sodium adducts were specific for their own cluster.

### 2.3. PRISM-M3C MS reveals sub-population-restricted itaconate and taurine changes in response to LPS treatment in microglia cells

Following validation of the PRISM MS with M3C methodology in GUVs, we applied this approach to LPS-treated microglial cells in search for candidate activation markers *in vitro*. Non-confluent populations of SIMA9 microglial-like cells were treated with LPS (0 to 500 ng/mL). Activation was confirmed by TNFα-sandwich ELISA and did not affect cell viability (**Figure S15 to S17**). In contrast but as expected, EOC microglial cells lacking toll like receptor 4 (TLR4) did not respond to LPS and served as negative control. After the DeepScan (20 µm) in negative ion mode, single cells were inferred from up to 7 connected pixels; larger areas of connected pixels were filtered out as likely cell aggregates, and candidate marker *m/z* were identified and annotated via METASPACE with FDR <=10%. Several *m/z* were upregulated after LPS treatment (**Figure 3a**). Most notably, the known microglial activation marker itaconate^39^ (*m/z* 129.02; [M-H]^-^) increased in a concentration-dependent manner (**Figure 3a,b** and **Figure S15)**. Of note, there are alternative annotations for *m/z* 129.02, *e.g.,* the structural itaconate isomers gluconate and mesaconate, in the KEGG database, which are unlikely in this biological context, but could not be ruled out by MS2 fragmentation (**Table S1 to S3; Dataset S1**). Interestingly, ion intensities (and presumed amounts of) itaconate in untreated cells were very low with low variance. In contrast, after LPS treatment some cells displayed very high intensities/levels (and many intermediate levels) of itaconate, whereas others stayed around zero, suggesting that the activation pattern was not homogenous for all cells and that subpopulations may exist including dead cells (**Figure 3b**). The *m/z* 124.01, most prominent in untreated, widely non-activated microglial cells and decreasing in response to LPS was identified as taurine, a known inhibitor of lysine demethylase activity and of microglial activation by LPS^40^ (**Figure 3b** and **Figure S18**). All *m/z* were validated using high-resolution FTICR MS and MS^2^ using SIRIUS (**Table S1 to S3; Dataset S1**)^41^.

**Figure 3.**
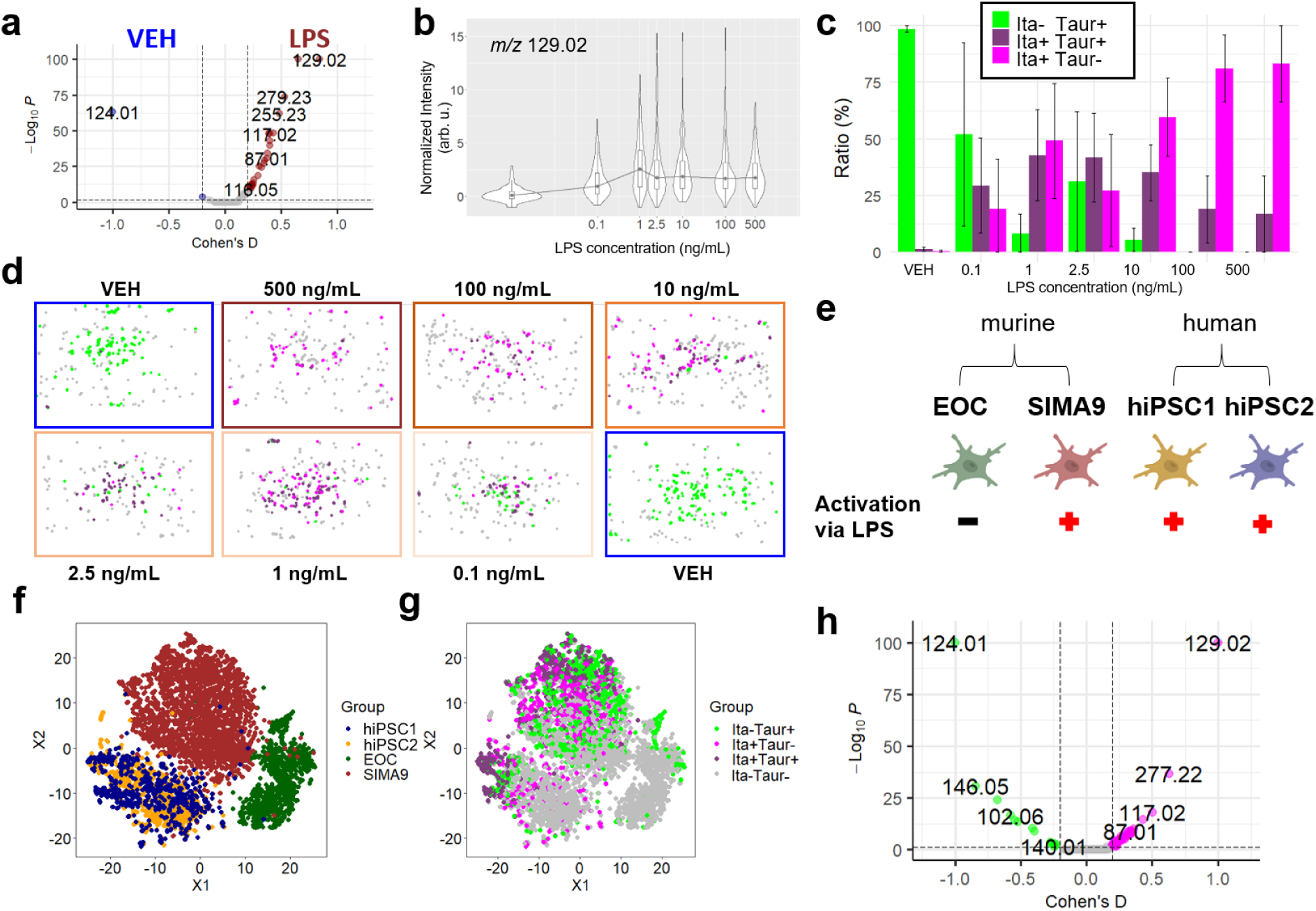
Microglial responses to LPS treatment revealed sub-populations of SIMA9 and hiPSC microglia cells. **a)** Modified Volcano plot for untreated (vehicle; VEH) and lipopolysaccharide (LPS) treated (500 ng/mL, 20h) SIMA9 cells highlights *m/z* 124.01 (taurine [M-H]^-^) and *m/z* 129.02 (itaconate [M-H]^-^) as markers for non-activated and LPS-activated microglial cells, respectively. **b)** Itaconate violin plot: normalized intensity of itaconate per cell indicates microglial activation in correlation with LPS treatment. **c)** M3C cluster analysis of SIMA9 cells for itaconate and taurine labeling. Cells are categorized into four distinct groups by post analysis – itaconate-positive (Ita+Taur-, pink), taurine-positive (Ita-Taur+, green) and positive for both markers (Ita+Taur+, purple). Cells that dońt fall into either category (Ita-Taur-, grey) are omitted. **d)** Visualization of M3C cluster analysis for chamber slide wells untreated (VEH) or treated with a concentration range of LPS (0.1 ng/mL to 500 ng/mL). **e)** Besides murine SIMA9 cells, the hiPSC-derived microglia cell lines hiPSC1 and hiPSC2 cells responded to LPS-treatment, as indicated by itaconate increases. Murine EOC cells lack toll-like receptor 4 and did not respond to LPS. **f)** t-distributed stochastic neighbor embedding (t-SNE) for the four microglial cells lines. **f)** t-SNE analysis of microglial activation status indicates that only a subset of SIMA9, hiPSC1 and hiPSC2 cells were activated by LPS (500 ng/mL for 20h): Cells were categorized as Ita-Taur+ (green), Ita+Taur+ (purple), Ita+Taur- (pink) or Ita-Taur- (gray). EOC cells were non-reactive. **h)** Volcano plot restricted to Ita-Taur+ and Ita+Taur-cells. Comparing both clusters yielded more defined *m/z*-signatures for microglial activation (15 markers of non-activation and 55 activation marker candidates) than using cell pools (2 markers of non-activation and 26 activation markers; compare a) by reducing extraneous data. In total 8.878 cells were analysed measured in six separate runs. This figure was partially created using BioRender.

We next applied M3C to PRISM-MS single-cell experiments. Using the activation-associated *m/z* for itaconate to check for cell activation, we noted k=2 subpopulations (but k=1 for EOC cells indicating no activation). k=2 was also found for taurine, the marker of non-activated microglia, which led to four possible labels for each single cell: taur+ita+, taur-ita+, taur+ita-, and taur-ita- (**Figure 3c,d**). The M3C approach can theoretically be extended to any (set of) *m/z* feature to identify subpopulations across all cells. However, performing this on a per-pixel level for MALDI imaging experiments, albeit feasible, would be very computationally intensive due to the large number of pixels considered. Using population statistics for the three of four subpopulations, we observed a shift from taurine-positive cells to itaconate-positive cells with increasing LPS concentration (**Figure 3c**). Taur-ita-cells (grey) were not considered for population statistics (**Figure 3d**). PRISM-MS with MC3 focusing on itaconate- and taurine-positive cells led to identification of 13 and 49 candidates for non-activation versus activation-specific biomarkers, respectively (**Figure S19**). For validation, we expanded the M3C clustering and used all activation-specific cell markers from **Figure 3a** instead of just itaconate, which generated a similar pattern to itaconate clustering (**Figure S20**).

Even though mouse SIMA9 cells have been used extensively as microglia surrogate as they display LPS responses^42,43^, their similarity with human microglia is rather limited. hiPSC-derived microglia may be a better model^11^. We therefore extended PRISM MS with M3C analysis to two hiPSC cell lines (hiPSC1and hiPSC2), which in contrast to EOC cells also displayed itaconate increases in response to LPS (**Figure 3e** and **Figure S21)**. As for SIMA9, k=2 resulted in the most stable clusters for the hiPSC lines and the same labels (taur+/ita+, taur-/ita+, taur+/ita- and taur-/ita-) were used. t-Distributed Stochastic Neighbor Embedding (t-SNE) analysis revealed a high degree of metabolomic similarity between both hiPSC cell lines and that EOC and SIMA9 cells were separate entities (**Figure 3f**). M3C-derived labels indicated that EOC cells were either non-activated or devoid of itaconate and taurine markers, whereas SIMA9 and the hiPSC lines were mixtures of cells belonging to any of the three subpopulations (taur+/ita+, taur-/ita+, taur+/ita-) (**Figure 3g**). M3C results for these three cell lines led to a refined taur-ita+ versus taur+ita-profile (**Figure 3h**; compare with Figure 3a) that disregarded unreactive and undefined cells in the analysis. Compared to the cell pool approach (**Figure 3a**), which identified 2 and 26 hits for non-activated versus LPS-treated cells, respectively, M3C analysis yielded 15 and 55 hits, respectively, from a total of 210 annotated m/z values - in addition to the information if a cell line was reactive to LPS or not (**Figure 3h**).

### 2.4 Translational single cell-informed MALDI imaging in organotypic hippocampal slice cultures suggests alterations in metabolic neuron-glia interplay in response to LPS-induced neuroinflammation

We next tested if marker candidates identified by single-cell analysis in mouse and hiPSCs translated well to endogenous microglia *in-vivo* or *ex-vivo*. To avoid LPS-injections *in-vivo*, we attempted to MALDI-image PRISM MS with M3C-derived micro-glial activation signatures in negative ion mode in LPS-treated and then cryosectioned organotypic rat hippocampal slice cultures, which feature ramified and widely non- activated microglia^44–48^. In these tissue cultures, microglial activation can be reliably studied in the presence of functional neuron networks and astrocyte syncytia^49^. Results can be compared with those gained in cell culture.

Slice cultures were split into three groups (**Figure 4a**): (1) Vehicle control (VEH) cultures were incubated for 72h in medium, (2) the LPS group slices were treated with 1 µg/mL LPS, and (3) the CLO group was exposed to 100 µg/mL clodronate liposomes. This treatment results in effective depletion of microglial cells by about 96% in slice cultures^46^. These cells engulf the liposomes that, in turn, release clodronate intracellularly, thus stopping the TCA cycle and causing apoptosis^44,45,50,51^. Five sets of slice cultures, with three conditions each, were subjected to MALDI-MSI. Comparison of Cohen’s D effect sizes between the VEH- and LPS-treated groups highlighted 8 and 11 annotations, respectively, as being specific for these tissues (**Figure 4b).** Itaconate was one of the markers specific for the LPS-treated slice cultures^31^ (**Figure 4c**). Immunofluorescence histology with anti-CD68 (red) and Hoechst stain (blue) performed after MSI identified CD68-positive cells only in LPS-treated tissue with apparent co-localization with a subset of itaconate-positive cells. As expected, itaconate was not observed in the VEH and CLO slices (**Figure 4c**).

**Figure 4.**
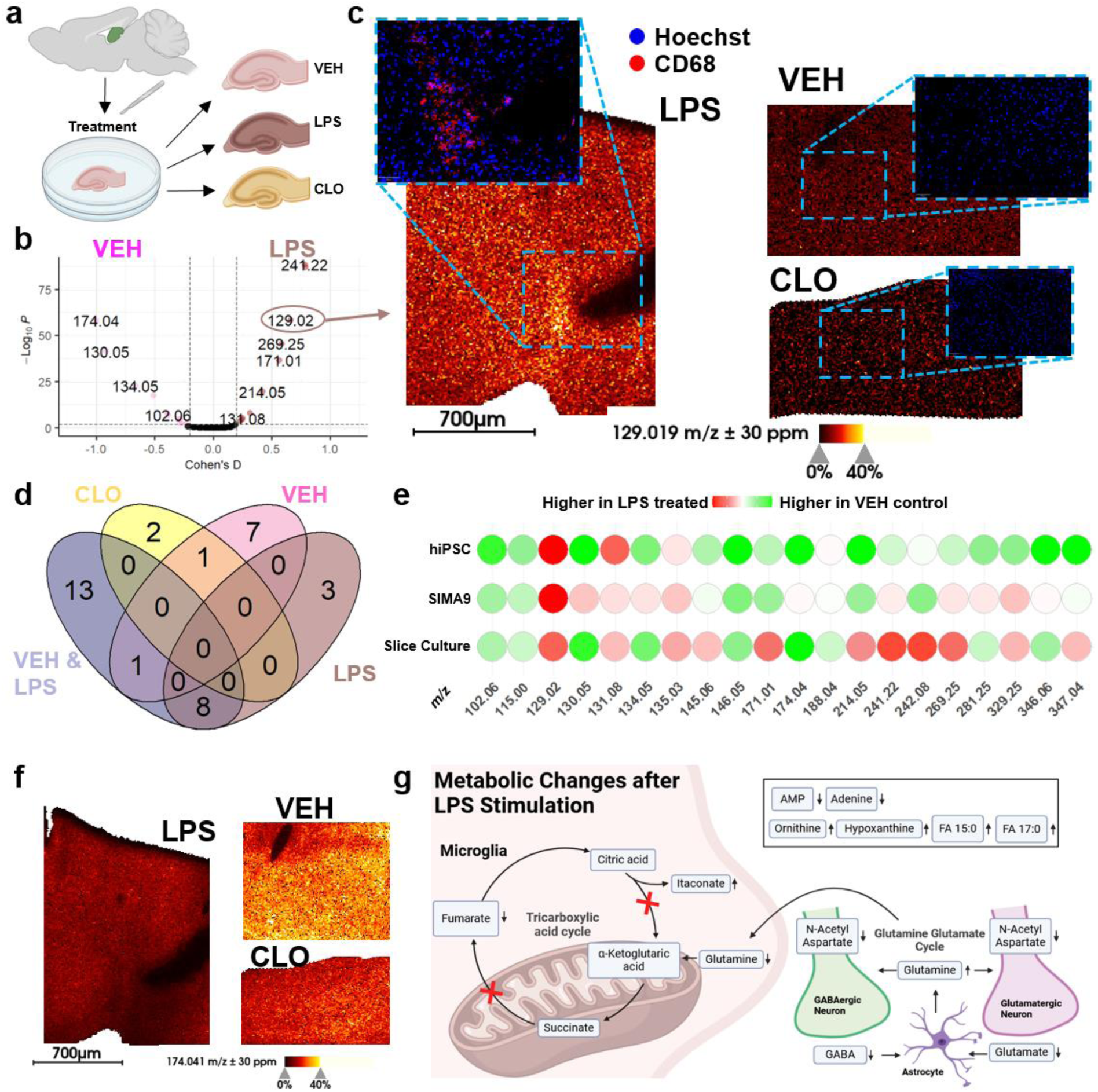
MALDI MS imaging of hippocampal slice cultures suggests metabolic neuron-glia interplay in response to LPS-induced neuroinflammation. **a)** Rat hippocampal slice cultures following 72h incubation with different treatments: vehicle-treated (VEH), 1 µg/mL lipopolysaccharide (LPS), or 100 µg/mL clodronate (CLO, a bisphosphonate for selective macrophage and microglia depletion)^44,45^. **b)** Volcano plot comparing VEH- and LPS-treated hippocampal slices revealed *m/z* features that mark non-activated and LPS-responding cells. Itaconate (*m/z* 129.02; [M-H]^-^) translates as a response marker from microglial cell populations (Figure 3) to slice cultures. **c)** Increased itaconate MSI ion intensities in LPS-treated hippocampal slices versus VEH and CLO controls (20 µm pixel size, negative ion mode). Anti-CD68-immunofluorescence overlay with Hoechst-stained nuclei suggests high microglia density in LPS-treated cultures. **d)** Metabolic profiling by MALDI MS imaging using KEGG METASPACE annotations at FDR<10%: Venn diagram comparing metabolites specific (Coheń s D>0.2 AND p<0.01) for CLO slice cultures versus VEH&LPS cultures containing microglia. Then comparing LPS-treated slice culture against VEH. **e)** Comparison of slice culture with hiPSC and SIMA9 cell metabolite profiles, suggesting common markers of non-activated microglia: GABA*(*m/z* 102.06), fumarate (*m/z* 115.00) and glutamate (*m/z* 146.05) and of active microglia: Itaconate (*m/z* 129.02), ornithine (*m/z* 131.08) and hypoxanthine (*m/z* 135.03). The hiPSC profile shared non-activated markers N-acetyl-alanine (*m/z* = 130.05), adenine (*m/z* = 134.05), N-acetyl-aspartate (NAA) (*m/z* 174.04), FA 18:1 (*m/z* 281.25) and AMP (*m/z* 346.06) with slice cultures. **f)** Ion intensity of NAA is reduced in LPS- treated slice culture compared to VEH and CLO tissue. **g)** Hypothetical model of metabolic neuron-glia interplay in LPS-activated hippocampal slice cultures. In total five separate measurements were performed, with all three conditions measured in triplicates each time, resulting in 45 slice culture sections. This figure was partially created using BioRender.

We compared microglia-containing slices (VEH and LPS) with microglia-depleted tissue (CLO) and identified 13 putative microglia metabolite markers, *e.g.*, *m/z* 191.02 (citrate/isocitrate) indicating an active TCA cycle (**Figure 4d**). 20 marker candidates from the VEH vs. LPS group comparison (**Figure 4b)** were used to generate an activation/non-activation microglia-specific profile that was then compared to the markers found in cell culture (**Figure 4e**). 11 out of 20 of the slice culture-derived activation marker candidates (all [M-H]^-^) showed the same trend (up or down after LPS-treatment) in hiPSC (9 out of 20 for SIMA9): *m/z* 102.06 (GABA), *m/z* 115.00 (fumarate), *m/z* 129.02 (itaconate), *m/z* 130.05 (N-acetyl-alanine), *m/z* 131.08 (ornithine), *m/z* 134.05 (adenine), *m/z* 135.03 (hypoxanthine), *m/z* 146.05 (glutamate), *m/z* 174.04 (N-acetyl-aspartate; NAA), *m/z* 281.25 (FA18:1 as general cell marker), *m/z* 346.06 (AMP). 6 out of 8 *m/z* features that shared a trend between SIMA9 and hiPSC translated to rat hippocampal slice cultures: *m/z* 102.06 (GABA), *m/z* 115.00 (fumarate), *m/z* 129.02 (itaconate), *m/z* 131.08 (ornithine), *m/z* 135.03 (hypoxanthine), *m/z* 146.05 (glutamate), but not *m/z* 171.01 (glycerol phosphate) and *m/z* 214.05 (glycerol-3-phosphoethanolamine; **Figure 4e**). All annotations were validated by accurate mass determination using magnetic resonance MS with <1 ppm mass accuracy and using MS2 fragment spectra-based annotation using SIRIUS^41^ (**Tables S1 to S3**). In contrast to larger metabolites like lipids where ion mobility differences of isobaric compounds can be exploited for imaging prm-PASEF-based fragmentation analysis, small metabolites <300 Da cannot be resolved well by ion mobility. Therefore, we employed QTOF mode fragmentation analysis. For MS2 identification of GABA other isomers (dimethylglycine, 3-aminoisobutyrate, 2-aminobutanoate or 3- aminoisobutanoate) cannot be ruled out. Because of the high brain concentrations and biological relevance of GABA (4-aminoisobutanoate), it is the most likely molecule though.

Interestingly, NAA (*m/z* 174.04), one of the markers that was higher in non-activated cells and VEH tissue was also higher in microglia-depleted CLO tissue (**Figure 4f**). NAA, a very abundant and clinical magnetic resonance spectroscopy-accessible brain metabolite that neurons store in large quantities^52,53^, is also modulated by LPS in microglia. For several metabolites like glutamine (*m/z* 145.06), the trends for LPS-induced changes (up/down) did not match: Glutamine was lower in activated cell populations compared with non-activated cells, while LPS-treated tissue showed higher overall glutamine levels compared to the control tissue. This apparent contradiction may, however, be explained by the fact that slice cultures represent functional networks of different neural cell types, whereas our single-cell analysis exclusively focused on microglia-like cells. It is tempting to speculate that co-localization of glutamine with itaconate in LPS-treated tissue (**Figure S22** and **23**) could suggest that activated microglial cells (marked by itaconate) accumulate glutamine from other cell types in their vicinity in tissue slices. Metabolic flexibility enables microglial cells to utilize glutamine as an energy source^54^.

This capability is underscored by the upregulation of glutaminase during neuro- inflammation, demonstrating their adaptation to use glutamine to meet energy demands^55,56^.The only source for glutamine in the brain is found in astrocytes, where glutamine synthetase converts glutamate to glutamine^57^. This conversion is part of the glutamine-glutamate cycle between neurons and astrocytes: The neurotransmitters GABA and glutamate are first taken up from the synaptic cleft by astrocytes and converted to glutamine. Then this glutamine is shuttled back to neurons to produce new neurotransmitters^58^. We also observed lower glutamate and GABA levels in slice and cell culture after LPS treatment, which may indicate that microglia that use glutamine as energy source may modulate the glutamine-glutamate cycle (**Figure S22** and **23**). To expand this hypothesis, reduced NAA levels in LPS-treated tissue could indicate that the glutamate reservoir in neurons is depleted to feed microglial glutamine needs whilst also trying to retain neurotransmitter balance (**Figure 4g**). In analogy with cell cultures, there was a trend towards lower taurine after LPS treatment in slice cultures, but (unlike cell cultures) this decrease was not significant. Assuming that no taurine precursors were provided through the slice culture medium, taurine may – similar to glutamine - simply be replenished by another cell type in hippocampal slices, most likely astrocytes^59^.

Our data suggests a hypothetical model of metabolic neuron-glia interplay in LPS-activated hippocampal slice cultures that can be more extensively tested: Microglial activation by LPS interferes with the TCA cycle by producing itaconate instead of α-ketoglutarate. Itaconate, in turn, inhibits succinate dehydrogenase and fumarate generation in the TCA cycle^30,31,60^. To replenish the TCA cycle, glutamine, predominantly produced by astrocytes, is utilized as alternative energy source by microglia. Therefore, the glutamine-glutamate cycle between astrocytes and glutamatergic/GABAergic neurons may be impaired, leading to lower levels of glutamate and GABA in LPS-treated slice cultures. The observed decrease in NAA within LPS-treated slice cultures could be a downstream effect, considering NAA’s exclusivity to neurons, alongside its role as a “glutamate reservoir” that facilitates the regeneration of glutamate and GABA (**Figure 4g**, **Figure 5, Figure S24**). Our results also support the notion that single cell (type) studies in culture need to be interpreted with caution, since compensatory metabolic and metabolite shuttling mechanisms might apply in multi-cell type tissues.

**Figure 5.**
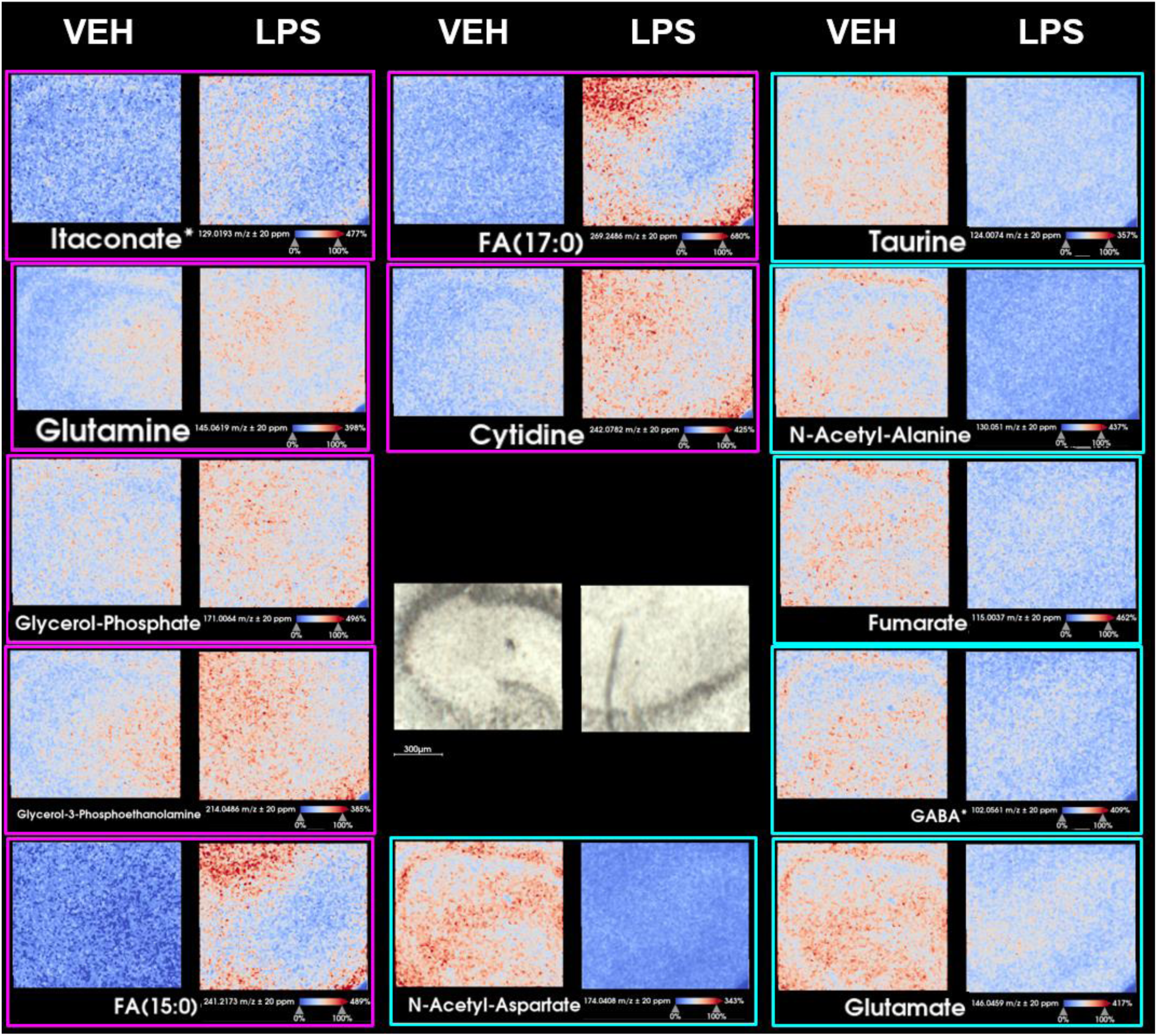
MALDI MS imaging reveals metabolic changes in hippocampal slice cultures. MALDI MS imaging of rat hippocampal slice cultures treated with vehicle (VEH) or with 1 µg/mL LPS at 5 µm pixel size. Ion images of metabolites that were significantly increased (magenta) or decreased (turquoise) between VEH and LPS groups (**Supplementary Tables S1 to S3**). For orientation, a bright field image of both tissue areas is included in the middle.

## 3. Conclusion

With PRISM-MS and M3C we enable fast and metabolite-preserving single-cell MSI of a wide range of lipids and hydrophilic, low mass (<200 Da) metabolites in unfixed cells. We introduce GUVs as a qualitative ground truth concept for MALDI MS imaging and present MSI of slice cultures as a means for providing biological context for single cell studies. PRISM-MS analysis of LPS-dependent microglia activiation highlights key changes of the itaconate-taurine ratio and of neuron-to-glia glutamate-glutamine shuttling. With its unique focus on small metabolites <200 Da that are fundamentally important for cell function and pathophysiology, PRISM-MS helps pave the way for single-cell low mass metabolomic analysis of cell sub-populations.

## 4. Methods

### 4.1. Materials

The MALDI matrix 1,5-diaminonaphthalene (DAN), poly-L-lysine, Tween-20, MTT reagent (thiazolyl blue formazan), trifluoroacetic acid (TFA), Mayer’s hemalaun solution, hydrochloric acid, sodium bicarbonate, magnesium sulfate, eosin Y-solution 0.5%, xylene, Biopore membranes and Eukitt were purchased from Merck KGaA (Darmstadt, Germany). Acetonitrile (ACN), ethanol (EtOH), hydroxypropyl methyl- cellulose, ammonium formate, polyvinyl pyrrolidone, and LC-MS grade water were from VWR Chemicals (Darmstadt, Germany). ESI-L low concentration tuning mix for calibration of the timsTOF flex mass spectrometer’s trap unit was used from Agilent Technologies (Waldbronn, Germany). Droplet-Micro-Array ITOs (DMA-ITOs) were acquired from Aquarray GmbH (Eggenstein-Leopoldshafen, Germany). Conductive indium tin oxide (ITO)-coated glass slides were sourced from Diamond Coatings (West Midlands, UK). BioGold microarray slides, Geltrex-coated surfaces, EDTA, DMEM/F12 with glutamine and HEPES, penicillin/streptomycin (PenStrep), Heat-inactivated fetal bovine serum (FBS), and heat inactivated horse serum, Dulbecco’s Balanced Salt Solution (DPBS), Hank’s Balanced Salt Solution, TrypLE Express, GlutaMAX supplement, 2-mercaptoethanol, and boronic acid were acquired from Thermo Fisher Scientific (Schwerte, Germany). From Sigma-Aldrich (Taufkirchen, Germany) L-Ascorbic acid 2-phosphate (LAAP), lipopolysaccharide (LPS) from Escherichia coli O55:B5, insulin, transferrin, pluronic acid coating solution, polyethyleneimine (PEI), laminin, MgCl_2_, CaCl_2_, progesterone, putrescine, and sodium selenite were obtained. Sucrose, 1,2-dioleoyl-sn-glycero-3-phosphocholine (DOPC), 1,2-dimyristoyl-sn-glycero-3-phosphocholine (DMPC), and 1,2-Dioleoyl-sn-glycero-3-phosphoetha- nolamin-N-(lissamin-rhodamin B-sulfonyl) (LissRhod-PE) were purchased from Avanti Polar Lipids (Alabaster, United States). Indium tin oxide (ITO) coated glass slides, for GUV preparation, were purchased from Visiontek Systems Ltd. (Chester, United Kingdom). FGF-2 (154), TGF-β1, (ROCK) inhibitor Y-27632, VEGF-165, SCF, M-CSF, IL-3, and GM-CSF were obtained from Cell Guidance Systems (Cambridge, UK). Sarstedt (Nümbrecht, Germany) provided 6-well plates and U-bottom 96-well plates. Silicone grease Rotisilon C/D was purchased from Carl Roth (Karlsruhe, Germany). Other materials included BMP4 and IL-34 from Miltenyi Biotec (Bergisch Gladbach, Germany), X-VIVO™15 from Lonza (Basel, Switzerland), and a 30 μm cell strainer from neoLab Migge GmbH (Heidelberg, Germany). Cell culture media included DMEM/F12, DMEM, PBS and EMEM from Capricorn Scientific (Ebsdorfergrund, Germany). 8-well chambers from IBIDI (Gräfelfing, Germany) were used. TNF-α levels were measured with a murine TNF-α ABTS ELISA Development Kit from PeproTech (ThermoFisher, Darmstadt, Germany). The ELISA buffer Kit was purchased from PeproTech. For slice cultures LPS from E. coli R515 was sourced from Enzo Life Sciences GmbH (Lörrach, Germany). Clodronate liposomes came from Liposoma B.V. (Amsterdam, Netherlands).

### 4.2. Cell lines and animal tissues

SIMA9 mouse microglia cells (Cat. No. CRL-3265), EOC 13.31 mouse microglia cells (Cat. No. CRL-2468), and LADMAC macrophages (Cat. No. CRL-2420) were obtained from ATCC (Manassas, United States). hiPSC lines hiPSC1 (male) and hiPSC1 (female) were used for microglial differentiation. Wistar rats were purchased from Janvier (Le Genest-Saint-Isle, France) and treated in accordance with the European directive 2010/63/EU and the ARRIVE guidelines. Consent of the animal welfare officers at the University of Heidelberg was obtained (license T37/21).

### 4.3. Generation, culture and LPS-treatment of hiPSC-derived microglia

All cells were maintained in humidified incubators at 37 °C and 5% CO_2_. hiPSCs were cultured as colonies on Geltrex-coated (15 mg/mL) 6-well plates. Geltrex was thawed on ice, diluted 1:50 in wash medium and subsequently added to tissue culture plates overnight at 4 °C for coating. hiPSCs were cultured in DMEM/F12 with glutamine and HEPES supplemented with Pen/Strep 1% (v/v), LAAP (64 µg/mL), sodium selenite (14 ng/mL), insulin 20 (µg/mL), transferrin 11 (µg/mL), FGF-2 100 (ng/mL), and TGF-β1 (2 ng/mL) with daily medium changes. Cells were passaged every 4-7 days at a 1:6 ratio using EDTA (0.5 mM) in DPBS. EDTA-containing liquid was aspirated, and hiPSCs were resuspended in hiPSC medium which was supplemented with the Rho-associated, coiled-coil containing protein kinase (ROCK) inhibitor Y-27632 for 24h after splitting to promote cell survival. Generation of hiPSC-derived microglia was based on published work with minor adaptations^61^. hiPSCs were dissociated to single cells with TrypLE Express, resuspended in wash media and centrifuged at 1200 x g for 3.5 min. Cells were resuspended in EB medium (hiPSC medium supplemented with BMP4 50 ng/mL, VEGF-165 (50 ng/mL), SCF (20 ng/mL), Y-27632 (20µM)) and plated into U-bottom 96-well plates at a density of 20,000 cells / well to form embryoid bodies (EBs). Cell numbers were determined using an automated Luna cell counter (Luna, Nümbrecht, Germany). Prior to cell seeding, U-bottom 96-well plates were coated with pluronic acid coating solution at RT for 15 min. A 50% media change was performed on day 2 and on day 4 EBs were replated into non-coated 6-well plates with 15 EBs / well using 1250 μL pipette tips (Sarstedt, Nümbrecht, Germany) that were manually cut to increase the diameter size and reduce the shear force on the EBs. All remaining EB medium was removed, and EBs were supplied with 3 mL / well macrophage progenitor medium X-VIVO™15 supplemented with Pen/Strep 1% (v/v), GlutaMAX supplement % (v/v), 2-mercaptoethanol (50 µM), M-CSF (100 ng/mL) and IL-3 (25 ng/mL). Medium was changed every 3 to 5 days depending on the consumption. Macrophage progenitor cells appeared in the supernatant after 3-4 weeks and could be harvested for many weeks, up to a few months. For maturation of microglia in monoculture, progenitor cells were harvested, filtered through a 30-μm cell strainer and plated on gold slides with IBIDI Chambers on top at a density of 4,000 cells / chamber. Prior to cell seeding, slides were coated with polyethyleneimine (1% v/v) and laminin (1 mg/mL). For coating, slides were first incubated with PEI coating solution (diluted 1:2000 in borate buffer (Boronic acid in H_2_O, adjust pH to 8.4 with NaOH, 25 mM)) for 10 min at RT. After washing 3 x with H_2_O, laminin (diluted 1:400 in DPBS + MgCl_2_ and CaCl_2_) was added overnight at 4 °C. IBIDI chambers were placed on top of the slides.

Macrophage progenitor cells were differentiated into mature microglia over a course of 1 week in macrophage maturation medium (Pen/Strep 1% (v/v), GlutaMAX supplement 1% (v/v), 2-mercaptoethanol 50 µM, N2-supplement (DMEM/F12 with glutamine) 70% (v/v), Pen/Strep 1% (v/v), insulin (500 µg/mL), progesterone (630 ng/mL), putrescine (1.611 mg/mL), sodium selenite (520 ng/mL), transferrin (10 mg/mL), IL-34 (100 ng/mL), GM-CSF (10 ng/mL)). Half of the IBIDI chambers per slide received a treatment with 500 ng/mL LPS in DPBS for 24h. Subsequently, the cell culture medium was aspirated, and the chambers were washed 3 x with ammonium formate buffer (150 mM) at 4 °C. This was followed by immediate snap-freezing of the slides in liquid nitrogen, ensuring that the slides did not come into direct contact with the liquid nitrogen by using an aluminum foil boat. IBIDI chambers were removed, and the slides were stored at -80 °C until further use.

### 4.4. Slice culture preparation, treatment and cryosectioning

Organotypic hippocampal slice cultures were prepared as reported previously^49^. In brief, hippocampal slices (400 µm) were cut with a McIlwain tissue chopper (Mickle Laboratory Engineering Company Ltd., Guildford, UK) from male rats at postnatal day nine (P9) under sterile conditions. No antibiotic was added. Three slices were placed on semipermeable Biopore™ membranes, at the interface between serum-containing culture medium and humidified normal atmosphere enriched with 5% CO_2_ (36.5 °C) in an incubator (Heracell, Thermo Fisher Scientific). The culture medium, exchanged three times per week, consisted of: 50% minimal essential medium (MEM), 25% Hank’s balanced salt solution, 25% heat-inactivated HS, glucose (4 mM) and L-glutamine (2 mM) at pH 7.3 (titrated with Trisbase). Slice cultures (group LPS) were stimulated at DIV 9 for 72 h to bacterial LPS (1 µg/mL) (from Escherichia coli, serotype R515 (Re)). Slice cultures (group CLO) were exposed for twelve days-in-vitro (DIV) to liposome-encapsulated clodronate to deplete the microglial cell population^46^. Liposomal clodronate was continuously present in the culture medium at a final concentration of 100 μg/ml from DIV 0 onwards^47,49^. Slice Cultures were snap-frozen on an aluminum block and stored at -80 °C. Prior to use, frozen slice cultures were placed in a cryostat (Leica CM1860 UV, Nussloch, Germany), where 15 µm cryosections were prepared and mounted onto ITO slides for measurement. The chamber temperature was set to -20 °C.

### 4.5. Giant unilamellar vesicle (GUV) preparation and slide preparation

GUVs were created by electroformation^62^ in a Vesicle Prep Pro device (Nanion Technologies GmbH, Munich). Lipids in chloroform were mixed at the desired ratios and 40 µL of 5 mM lipid mix was spread onto the conductive side of an ITO (indium tin oxide)- coated slide. The slide was left under vacuum in a desiccator for 30 min to evaporate the chloroform. Afterwards, a rubber ring was covered in silicone grease and positioned onto the spread lipids. 275 µL of sucrose solution (320 mM) were heated to 60 °C and filled inside the ring. A second ITO slide was placed on top of the ring with the conductive side facing down and the slides were connected to the respective electrodes inside the Vesicle Prep Pro. The Standard program that applies an AC field (3 V, 5 Hz) for 138 min was run at different temperatures. For the lipid mix containing 99% DOPC and 1% LissRhod-PE, the temperature was set to 37 °C and the GUVs were collected immediately after. For the other lipid mixes (99% DMPC, 1% LissRhod-PE and 49.5% DOPC, 49.5% DMPC, 1% LissRhod-PE), it was set to 70 °C and the GUVs were collected after 5 h when the chamber reached room temperature (RT) and slowly cooled to 12 °C in a thermoshaker (MKR 23, Hettich Benelux, Geldermalsen) over 1 h. All GUVs were eventually stored at 4 °C. GUVs were diluted 1:2000 in sucrose and pipetted onto ITO-Slides. These slides were coated using 4% BSA beforehand. These were then imaged with an LMD7000 laser microdissection microscope (Leica Microsystems, Wetzlar, Germany), where fluorescence signals were measured using the 515-545 nm emission filter channel at 10x magnification.

### 4.6. PRISM-MS MALDI imaging

#### 4.6.1. Slide preparation for single-cell MALDI-MS imaging metabolomics

Gold-coated microarray slides were initially cleaned by sonication in three different solutions: acetone, methanol, and ddH_2_O, each for 10 min, followed by air-drying in a desiccator. For calibration, teach marks were applied to the slides, which were then scanned using a CanoScan 8800F (Canon, Tokyo, Japan) slide scanner operated with Silverfast software. The slides were sterilized using 80% ethanol and dried under a sterile bench. They were then coated with sterile 0.01% (w/v) poly-L-lysine for 5 min at RT. After coating, slides were washed thrice with sterile ddH_2_O and dried again under sterile conditions for 2h.

#### 4.6.2. Sample Preparation

Frozen slides were inserted into a lyophilization device, the Alpha 1-2 LDplus (Christ, Osterode, Germany), and dried for 60 min. Immediately following this drying step, for the PRISM-MS workflow, the matrix was directly applied by spray-coating. To mimic published optically guided workflows for comparative analysis with the PRISM-MS workflow, a second slide was positioned in a sterile hood at room temperature for 30 min prior to matrix spray-coating.

#### 4.6.3. Matrix spray-coating

1,5-Diaminonaphthalene (DAN) matrix solution (10 mg/mL) was prepared in ACN/ H_2_O/Trp-D5 (60:39:1, v/v). Trp-D5 (from 5 mg/mL stock) was used as internal standard. Matrix solutions were sonicated for 15 min and deposited in eight spraying cycles onto slides using an M5 Sprayer (HTX Technologies LLC, Chapel Hill, USA). The spraying parameters were as follows: solvent composition was 60% ACN in water; nozzle temperature 70 °C; and bed temperature at 35 °C; flow rate at 0.07 mL/min with a nozzle velocity of 1200 mm/min. The track spacing was maintained at 2 mm in a HH pattern. The spraying pressure was 10 psi, and the gas flow rate was 2 L/min. There was no dry time involved in the process. The height of the nozzle from the slide surface was set at 40 mm. Following matrix-coating, slides were immediately placed in the mass spectrometer.

#### 4.6.4 PRISM MS PreScan

PreScans at 200 µm pixel size were performed using the methods listed using an orthogonal TimsTOF flex mass spectrometer (Bruker Daltonics) in QTOF mode with flexImaging 7.4 software (Bruker Daltonics).

| Parameter | PreScan (Cells or Tissue) | DeepScan (Cells or Tissue) | PreScan (GUVs) | DeepScan (GUVs) |
| --- | --- | --- | --- | --- |
| Polarity | negative | negative | positive | positive |
| Range [$m/z$] | 50-700 | 50-700 | 400-950 | 400-950 |
| Laser shots | 20 | 150 | 20 | 200 |
| Laser frequency [1/s] | 10,000 | 10,000 | 10,000 | 10,000 |
| Laser field size [$\mu\text{m}$] | 200x200 | 20x20 | 200x200 | 20x20 |
| Funnel 1 RF [Vpp] | 150 | 150 | 420 | 420 |
| Funnel 2 RF [Vpp] | 160 | 160 | 400 | 400 |
| Multipole RF [Vpp] | 170 | 170 | 380 | 380 |
| Collision Energy [eV] | 3 | 3 | 10 | 10 |
| Collision RF [Vpp] | 500 | 500 | 1800 | 1800 |
| Low $m/z$ | 70 | 70 | 300 | 300 |
| Transfer Time [$\mu\text{s}$] | 40 | 40 | 85 | 85 |
| Pre Pulse Storage [$\mu\text{s}$] | 4 | 4 | 10 | 10 |

For tissue and cell measurements, the instrument was calibrated using ESI tune mix as calibrant and single point online calibration using a matrix peak of 1,5-DAN [M-H]^-^ at *m/z* 157.0771. For measurements of GUVs in positive ion mode, red phosphorous was used as calibrant without additional online calibration. The laser was set to custom for PreScans with smart beam set to M5 and Beam Scan to on. The scan range was set to 126 resulting in a field size of 200 µm. The 20 µm DeepScans were performed using default 20-micron imaging laser settings. MS2 spectra were acquired using the same method as for the DeepScan with an *m/z* selection window of 0.5 *m/z* and collision energies varying between 5-30V. The resolution was 30,000 at *m/z* 208.1140.

#### 4.6.5 PRISM-MS data processing, data storage and DeepScan

The timsTOF flex MS imaging .tdf files were converted into .imzML files through *timsconvert* with a precision of 32 bit, excluding ion mobility and without compression^63^. The .imzML and corresponding .mis files were then analyzed by the following PRISM- MS R-script:

(i) PreScan imaging data in .imzML format is loaded with a tolerance of 10 ppm using the Cardinal open-source software^64^. Additionally, the .mis-file created by the PreScan is loaded using the R-package *xml2* (https://github.com/r-lib/xml2) to parse the file. The script identifies predefined molecular peaks, such as the 281.2485 *m/z* peak corresponding to ubiquitous FA 18:1 used for targeting in single cells. (ii) A threshold κ defined as

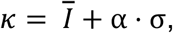

where *Ī* denotes the mean intensity of a given *m/z* feature per ion image, σ the standard deviation and α being an integer value, is then applied to these peaks to generate a mask that highlights pixels where the peak is present, thus indicating areas of interest within the sample. The threshold can be adjusted manually, but defaults is set to α = 1.

(iii) This mask is converted into a binary image aligned with the original flexImaging coordinates, thus allowing for definition of regions of interest (ROIs). It integrates the .mis file information with the .imzML data coordinates, and the binary image is resized to match the resolution specified in the .mis file. An optional ROI dilation step is included, set by default to 1, indicating no change. (iv) Contours of these ROIs are derived from the thresholded image and then used as ROI coordinates for the subsequent DeepScan. Each contour defines a new imaging region, which is recorded in a newly generated .mis file. This new .mis file is a modification of the original, updating the measurement method name and incorporating the new raster size and coordinates for each new ROI. (v) This file can then be directly loaded into flexImaging for targeted data acquisition within the specified ROI.

MSI data was converted to .imzML-files again, which were then uploaded to the false discovery rate (FDR-) controlled Metaspace database for annotation using HMDB (endogenous) and KEGG pathway databases^37^. Annotations with FDR ≤10% were then exported as a .csv file to store all annotations. An in-house R-script reduced the extracted .imzMLs to the annotated peaks with a tolerance of +/- 0.005 Da with the help of the Cardinal R package for data storage until further processing^64^.

### 4.7. On-tissue timsTOF MS2 data analysis

MS2 fragmentation spectra files were opened using DataAnalysis (Bruker Daltonik) and directly exported as .mgf files. These files where then loaded into and evaluated with SIRIUS Version 5.8.6^41^.

### 4.8. MALDI magnetic resonance MSI (MR-MSI)

Ultra-high resolving power data was acquired on a solariX 7T XR Fourier Transform Ion Cyclotron Resonance (FT-ICR) mass spectrometer (Bruker Daltonics), equipped with a smartbeam II 2 kHz laser and ftms control 2.3.0 software (Bruker Daltonics, Build 92). Mass spectra were acquired in negative ion mode (*m/z* range 71.66–1000) and a time domain of acquisition of 8 M, resulting in long free induction decay (FID) times of 1.39 s and a mass resolution of 700,000 at *m/z* 208.1140. Ion optics settings were constant for all measurements: funnel RF amplitude (1000 Vpp), source octopol (5 MHz, 350 Vpp), and collision cell voltage: 0 V, cell: 2 MHz, 1000 Vpp). The source DC optics was also constant for all measurements (capillary exit: -150 V, deflector plate: -200 V, funnel 1: -150 V, skimmer 1: -15 V), as well as the ParaCell parameters (transfer exit lens: 20 V, analyzer entrance: 10 V, sidekick: 0 V, side kick offset: 1.5 V, front/back trap plate: -2.8 V, back trap plate quench: 30 V). Sweep excitation power for ion detection was set to 14 %, and ion accumulation time was 0.05 s. For IMP and ISOM, the transfer optics were as follows: time of flight: 0.4 ms, frequency: 6 MHz, and RF amplitude: 350 Vpp. The laser parameters were laser power: 30 %, laser shots: 150, laser frequency: 2000 Hz, and laser focus: medium, at a lateral step size of 40 µm.

### 4.9. Single cell and GUV data analysis

Annotated and peak-picked .imzML files for single cell analysis were processed using an in-house R script, employing the Cardinal package for loading mass spectrometry (MS) data^64^: (i) The script first extracts labels, constructs an intensity matrix, and stores essential image metadata such as pixel coordinates. (ii) Data normalization is performed against the internal standard (IS) Trp-D5 (*m/z* 208.1140) by dividing the ion intensity at each point in a matrix ***M*** ∈ ℝ^*i*×*j*^by the Trp-D5 intensity in the same spectrum. Here, each row *i* in the intensity matrix ***M*** corresponds to a distinct spectrum and each column *j* to unique *m/z* values. The normalization results in a new matrix ***M***^′^, representing normalized intensity values:

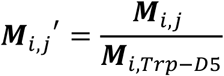

(iii) After normalization, Z-scale standardization is implemented for each *m/z* column. This standardization ensures a consistent scale across the entire dataset by adjusting the normalized intensity values. Specifically, each value is modified such that the distribution of intensities for each *m/z* feature has an average μ_*j*_ of zero and a standard deviation (σ_*j*_ of one. A new matrix ***M***^′′^ represents normalized, scaled intensities:

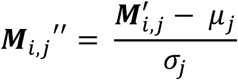

This two-step process corrects for potential discrepancies arising from sample handling, instrument performance, and other factors. Consequently, it enables more reliable comparison of molecule abundances across spectra. (iv) A cell-selection algorithm in the script then detects cell signal peaks such as *m/z* 281.2485 (or any other *m/z* chosen by the user). (v) Pixels with normalized and scaled intensities ≤3, for this feature, are removed from the imaging dataset. (vi) The remaining pixels are then processed through an algorithm that identifies and clusters connected pixels as cells, excluding clusters larger than 20 pixels. To focus on true single cells, we showed that setting this to seven pixels excludes larger cell clusters. The script calculates and stores the mean spectrum for each cell cluster. This process is applied to all single cell measurements. For Giant Unilamellar Vesicles (GUV) processing, data underwent Total Ion Current (TIC) normalization, and the GUV-specific signals were the lipids DOPC [M+H]+ (*m/z* 786.6) and DMPC [M+H]+ (*m/z* 678.5). Here masks from both peaks are added together to pick GUVs, since there are no common peaks for both GUVs. Slice culture data was also normalized against Trp-D5 *m/z* 208.1140, then z-scale standardized and stored in a data frame for further processing.

### 4.10. Monte Carlo consensus clustering (M3C)

We used Monte Carlo consensus clustering (M3C) for analysis of cluster stability and for determination of the optimal number of clusters (K) for a given *m/z*-feature in a cell population^38^. This approach utilizes hierarchical clustering, conducted over five iterations per dataset on a single CPU core, ensuring methodological consistency. To guarantee reproducibility, a fixed seed value of 42 was used across all analyses. We restricted the analysis to a maximum of eight clusters (maxK) and executed 200 real and 200 reference iterations per cluster number (k) to evaluate the Relative Cluster Stability Indices (RCSI) and to calculate p-values. The RCSI metric was used to measure the stability of clustering outcomes, reflecting the consistency of sample groupings across multiple iterations; higher RCSI values indicate greater clustering stability. The significance of each cluster number was assessed based on p-values, with a threshold set at less than 0.01 indicating statistically significant clustering distinct from random chance. The optimal k was selected based on the highest RCSI among all k values with a p-value lower than 0.01. If no k values met this significance criterion, it was concluded that the observed *m/z*-feature did not exhibit sub clustering, thus suggesting a homogeneous distribution across the studied population.

### 4.11. Volcano Plots using Coheńs D

Volcano plots were generated using in house R-code, which is included in the available processing script. For every feature of the data, i.e. for a given *m/z*-interval, a p-value was generated that was adjusted according to the Benjamini Hochberg criterion to account for high dimensionality of the data^65^. Coheńs D effect size was calculated for each group using this script. Since the data was standardized, the standard deviation σ_12_ is 1 and μ_1_ and μ_2_ are the mean intensities for a given group.

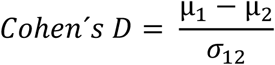

A peak was considered relevant, if the effect size was above +0.2 or below -0.2^66^ and the corresponding adjusted p-value was below 0.05. For cell culture, all data points were chosen, since they were considered being independent cells. In the slice culture analysis, a sample of 1,000 pixels was randomly selected for each condition from a total population of 1,284,814 pixels. This random selection process was repeated 10 times. Subsequently, the mean p-values and mean effect sizes were calculated based on these statistical samples. This procedure was implemented to eliminate the statistical dependence between adjacent pixels, which could lead to artificially inflated p-values.

## Data availability statement

Extensive data (MR-MSI data; TIMS-MSI-MS and MS^2^ data incl. fragment spectra and images) is provided as **Supplementary Datasets**. All data will be made publicly available in line with requirements by the accepting journal.

## Ethics approval statement

hiPSCs used in this study were generated from healthy donors. Donors gave written informed consent. All experiments with human material are in accordance with the declaration of Helsinki.

## Code availability

Code will be made publicly available upon tentative acceptance of the manuscript in line with requirements by the accepting journal.

## Acknowledgements

The authors thank Thomas Enzlein and Denis Abu Sammour for helpful discussions and insightful comments on the manuscript.

## Author contributions

J.L.C. conceived the PRISM-MS concept, wrote all code, performed MSI experiments, analyzed MSI data, prepared Figures and contributed to the first draft of the manuscript. J.H. performed cell culture, LPS-stimulation and MSI experiments, ELISA, and she analyzed MSI data. T.B. performed MS2 experiments. S.S. suggested experiments and analyzed data. A.L. and O.K. conduced hippocampal slice culture experiments and compound treatment. S.M. and K.G. generated and characterized GUVs. J.J. and P.K. generated, propagated and analyzed hiPSCs. P.L. generated droplet microarrays. D.S. and C.H. conceived the study and discussed results. C.H. provided infrastructure, supervised the overall study, conceived GUVs as a ground truth in MSI, suggested experiments and wrote the final manuscript. All co-authors contributed to and edited the manuscript.

