## Supporting Information for "Fast Single-Cell MALDI Imaging of Low-Mass Metabolites Reveals Cellular Activation Markers"

### Supplementary Methods

##### Cultivation of murine cell lines and LPS-treatment

SIMA9 cells were grown in DMEM/F12 medium, supplemented with 1% PenStrep, 10% (v/v) FBS, and 5% (v/v) HS. EOC13.31 cells were cultured in DMEM medium, supplemented with 1% PenStrep, 10% (v/v) FBS and 20% LADMAC macrophage-conditioned media. Cells were maintained at 37 °C in a humidified incubator containing 5% $CO_2$. Sub-culturing was performed every second day at a ratio of 1:5 or 1:2 for SIMA9 cells or EOC13.31, respectively. Both cell lines were confirmed to be mycoplasma-free at the last passage. For each experiment, 1000 cells/well (5 cells/µL) were seeded into 8-well IBIDI chambers on BioGold gold-coated slides and incubated for 24h. For LPS treatment, the media was replaced with serum-free media, and cells were treated with PBS as vehicle control (VEH) or stimulated with LPS at various concentrations (0.1, 1, 2.5, 10, 100, or 500 ng/mL) for 20h. Cell culture supernatants were stored at -80 °C for subsequent ELISA analysis. For single-cell MALDI MS imaging, the slides were cooled on ice and washed thrice with ice-cold 150 mM ammonium formate. The slides were then snap-frozen in liquid nitrogen, the IBIDI chamber was detached, and the slides were then stored at -80 °C until further use.

##### Cell viability assay

Cells were seeded and treated (VEH or LPS) as described above. Fresh MTT stock (5 mg/mL) was prepared in PBS, sonicated for 10 min, and sterile-filtered. For the MTT assay, half of the culture media was removed from each well, and 10 µL MTT was added to the remaining volume, followed by incubation for 4h at 37 °C. Subsequently, 100 µL of 10% SDS in 0.01M HCl was added to each well and incubated overnight at the same temperature. Absorbance was measured at 570 nm and 670 nm using a POLARstar (BMG Labtech, Ortenberg). Data analysis involved normalization to VEH and statistical testing using one-way ANOVA. ****$p < 0.0001$, **$p < 0.01$, *$p < 0.05$ to Vehicle (VEH; 0 ng/mL).

#### ELISA

TNF-α measurement in cell culture supernatants utilized a sandwich ELISA according to manufacturer's instructions. Color development was monitored at 5 min intervals for 5-40 min with a spectrophotometer (Multiscan Spectrum, Thermo Fisher Scientific) at 405 nm with wavelength correction set at 650 nm. TNF-α concentrations were calculated based on the calibration curve. Data analysis was done using an in house R-script, statistical testing employed one-way ANOVA ****$P < 0.0001$, **$P < 0.01$, * $P < 0.05$ to Vehicle (VEH; 0 ng/mL).

### Supplementary Figures

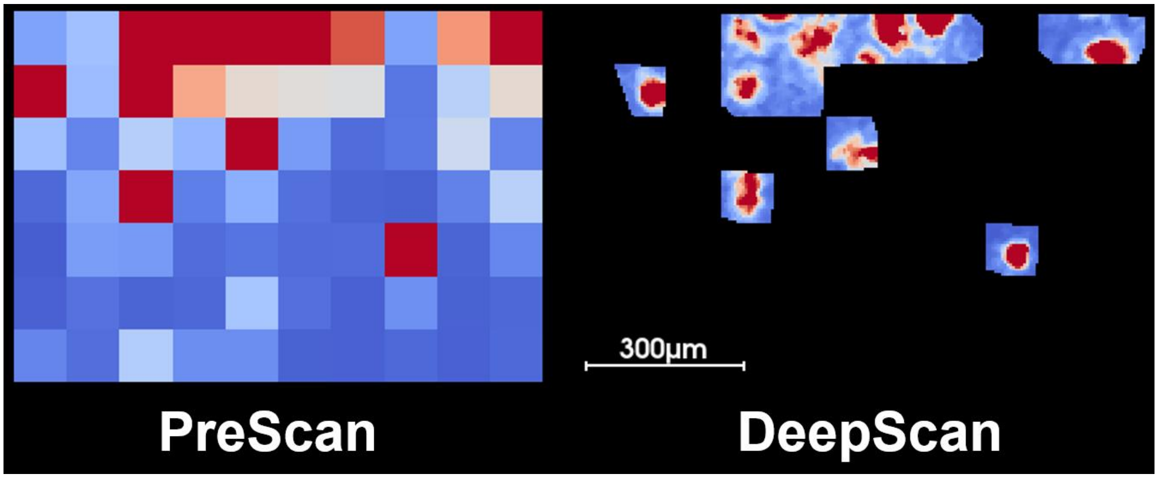

Figure S1. PRISM-MS workflow for SIMA9 Cells using 100 µm PreScan and 5 µm DeepScan. The detection feature for generating the mask was FA(18:1)[M-H]$^-$ with *m/z* 281.25, a spot dilation factor of 1 was used and the threshold was set to be the mean intensity + one standard deviation (σ).

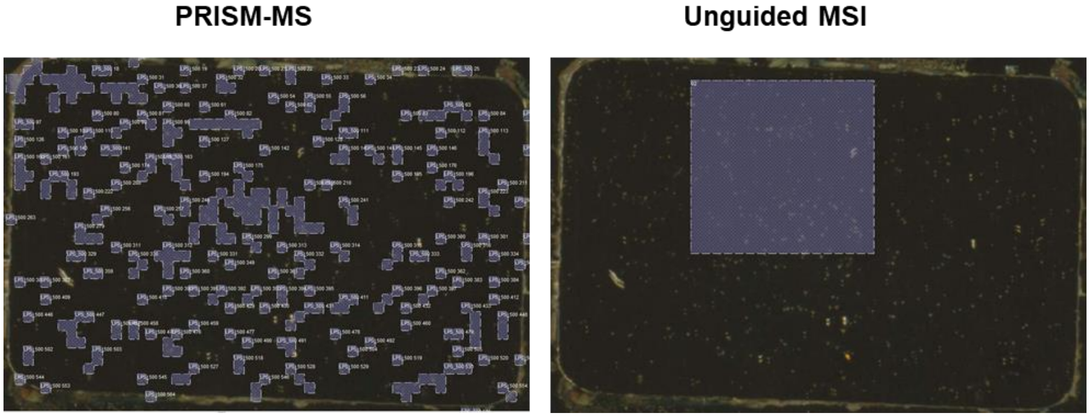

Figure S2. Comparison of “Cell” and Area Coverage using PRISM MS vs. Unguided Whole-slide MSI. PRISM MS versus unguided MSI with equal pixel counts on the same sample (SIMA9 cells seeded on a gold slide). The image was obtained with a scanner (CanoScan 8800F, Canon), and cells were visible as golden dots. Left: Mask of “cell”-containing pixels using the PRISM-MS PreScan with a threshold of mean intensity plus one s.d. (σ) of *m/z* 281.25 (FA18:1[M-H]$^-$) and a spot dilation of 1. Right: Unguided MSI covering identical total area with similar measurement time. Note that PRISM MS captures more “cells” and a greater area, while simultaneously reducing the volume of unnecessary data. Both images capture areas of 40,000 pixels at 20 µm pixel size.

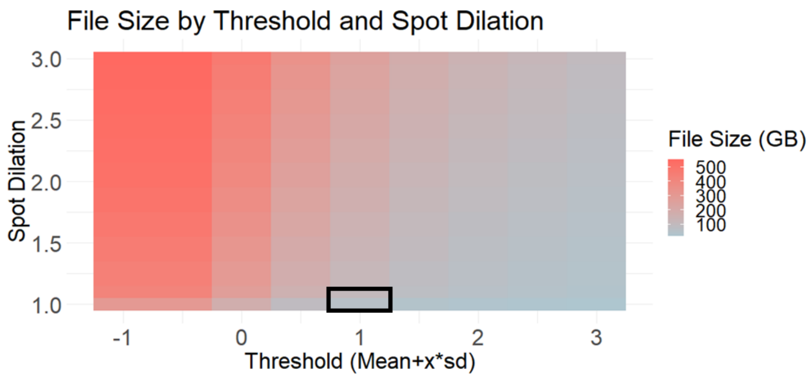

Figure S3. Effect of Spot Dilation and Thresholding on File Size. Illustrates the impact of spot dilation and thresholding on file sizes, for a DeepScan at 20 µm pixel size, using *m/z* 281.2485 (FA18:1) as PreScan marker. Applying a spot dilation of 1 and a threshold of mean intensity + one standard deviation (σ) (black box) yielded a file size of 50 GB for a complete slide. Scanning the complete slide at 20 µm, using the same settings, would have resulted in a file larger than 500 GB. Since file size is directly proportional to measurement duration, this also illustrates the time saved using PRISM-MS, besides reducing the amount of unwanted data. Using higher thresholds can reduce data size/acquisition time even further, with the trade-off being that not all "cells" are capturing.

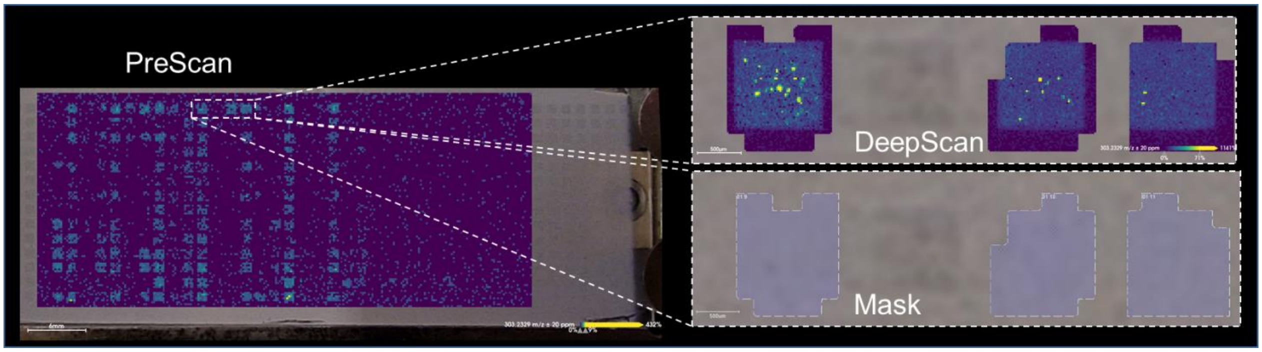

Figure S4. PRISM-MS on a 14 x 48 DMA-ITO Slide with SIMA9 cells. Left: PreScan visualizing *m/z* 303.23 (arachidonic acid; $[M-H]^-$) as marker for SIMA9 microglial cells cultured on a 672-spot Droplet-Microarray (DMA). Presence of cells is represented via the viridis color scale. PreScan region selection was performed with a threshold of the mean *m/z* 303.23 intensity + one s.d. (σ) and a spot dilation of 1.25 to create the mask for the subsequent DeepScan. Lower right: Mask for the DeepScan. Top right: DeepScan displaying arachidonic acid as cell marker (*m/z* = 303.23) at 20 µm pixel size.

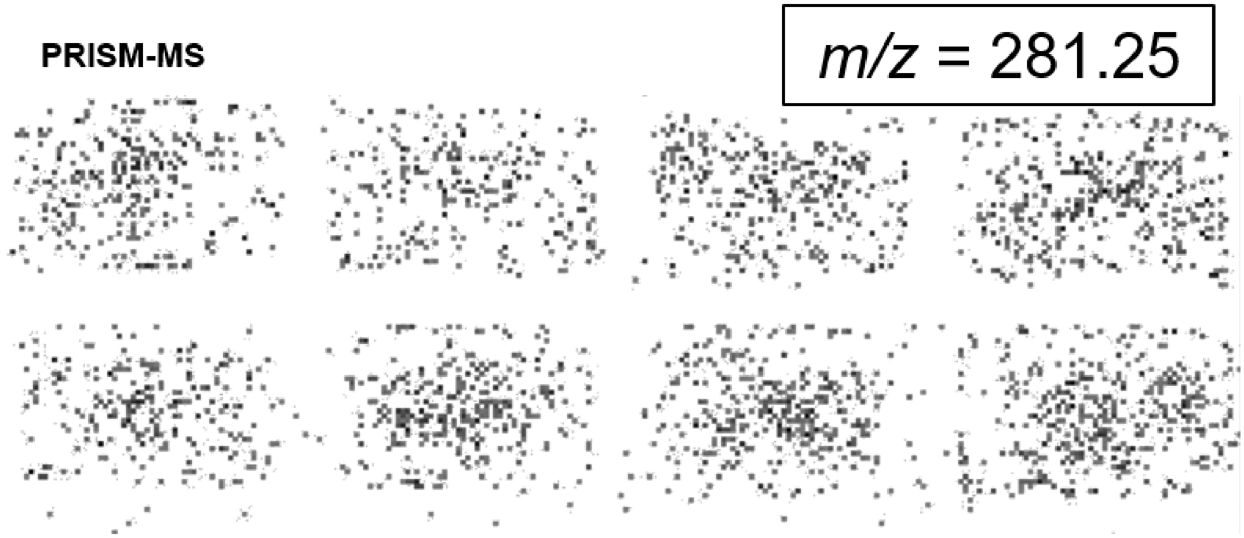

Figure S5. PRISM-MS PreScan: Distribution of SIMA9 Microglial Cells on Ibidi Chamber Slides. The mask created after a PreScan using the fatty acid FA (18:1) (*m/z* = 281.25) as marker for SIMA9 microglial cells cultured on 8-well Ibidi chamber slides. Spot dilation was 1 and threshold mean intensity + one s.d. (σ). Presence (grey pixels) or absence (white pixels) of cells is indicated.

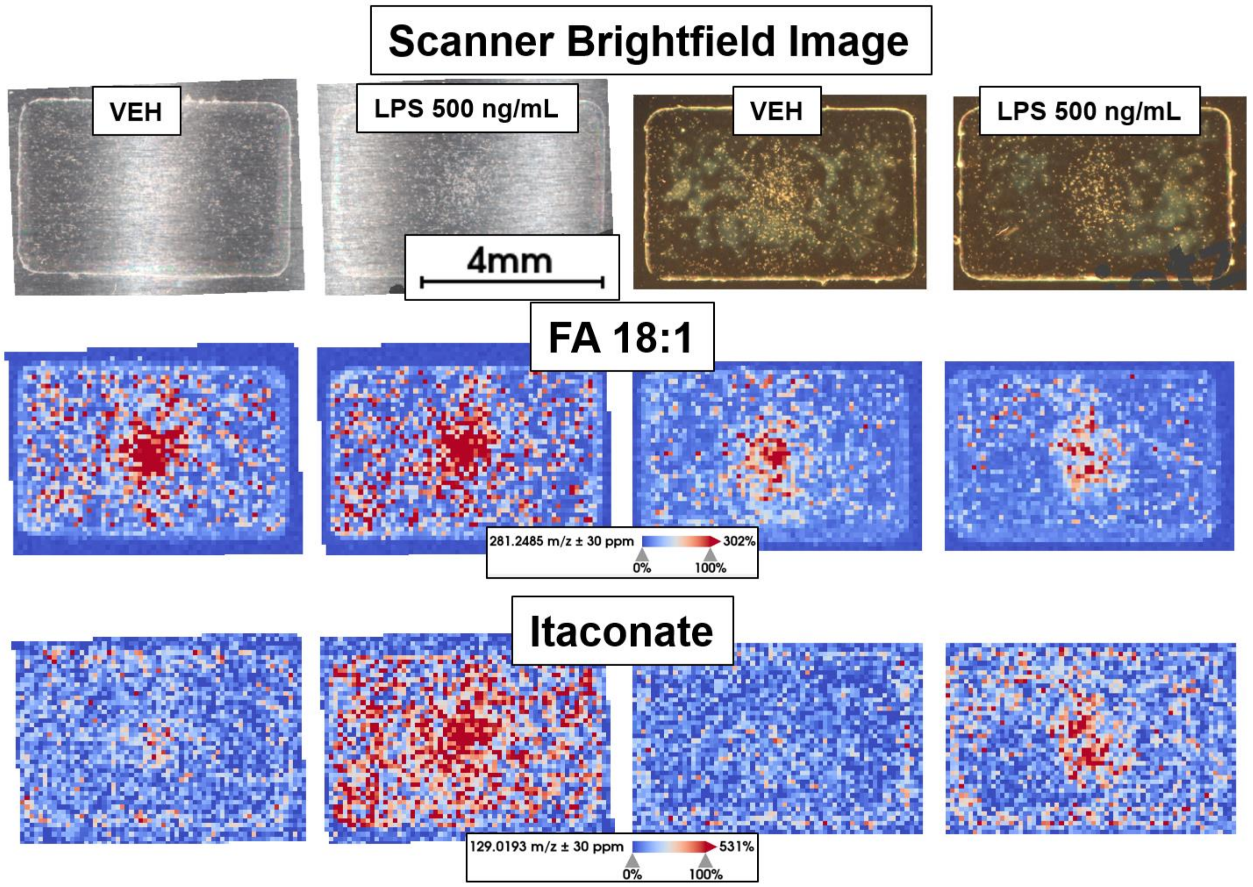

Figure S6. PRISM-MS can Measure on Gold and Steel Slides. SIMA9 cells were seeded on steel and gold slides in VEH, PBS-treated, and 500 ng/mL LPS-treated conditions. A bright field scanner (CanoScan 8800F) captured an overview image to roughly estimate cell presence, but cell detection at this level was not possible. This demonstrated that PRISM-MS enables MALDI-MSI on non-transparent surfaces and that cells can be cultured on these surfaces. Further research is needed to compare cell growth on gold, metal, and ITO slides for MSI. A PreScan was conducted on two slides at 200 µm pixel size. No DeepScan was performed in this proof of concept experiment. Spectra were uploaded to Metaspace, showing 64 KEGG annotations at FDR=10% for the metal slide and 40 for the gold slide.

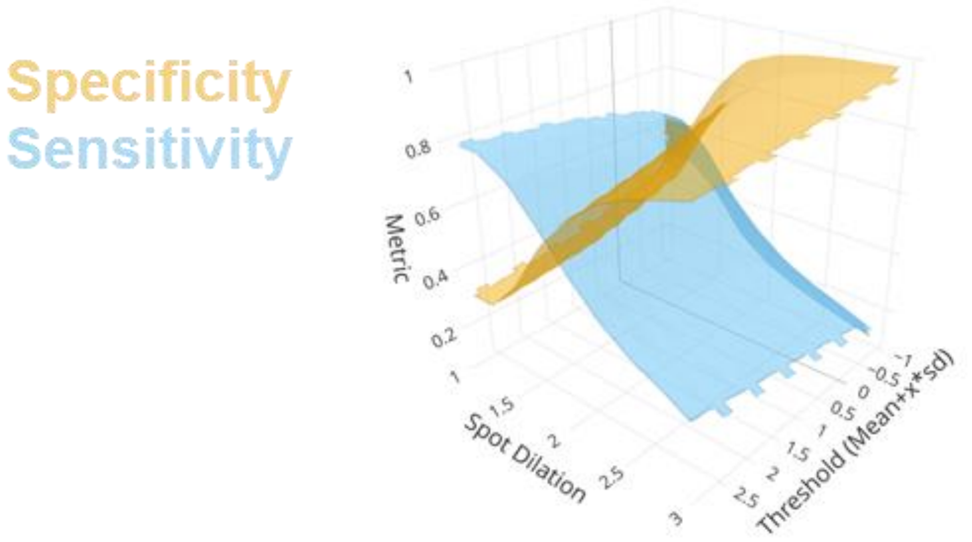

Figure S7. Impact of Spot Dilations and Thresholds on Specificity and Sensitivity of PreScan for Detection of SIMA9 Microglia Cells.

The ground truth of positive cells was established by Hoechst staining of SIMA9 cells and fluorescence scanning. Employing *m/z* = 281.25 (FA18:1) as the defining feature for “cells”, specificity and sensitivity refer to the ability to correctly identify cells whilst avoiding measuring background. Specificity measures the proportion of true negatives (*i.e.,* correctly identified background) out of all pixels identified as negatives. Sensitivity measures the proportion of true positives (*i.e.,* correctly identified cells) out of all actual positives. The metric used here is the percentage of Hoechst-positive pixels/”cells” that was correctly identified using m/z 281.25 intensity. It was obtained by analyzing the pixel overlay from the Hoechst-stained fluorescence image and a MALDI MSI PreScan at a 200-µm pixel size. At this pixel size, the analysis checks how much of the Hoechst-stained area is identified with different PreScan parameters. Different spot dilation factors (1 to 3) that expand the area deemed “cell”-positive in the PreScan as a safe-guarding measure prior to the DeepScan were evaluated. Moreover, various intensity thresholds for mean intensity of *m/z* = 281.25 and -1 to +2.5 s.d. were tested. Setting this threshold as well as spot dilation depends on the desired outcome and experiment. In this study we mostly used a dilation factor of 1 to 1.5 and a threshold of mean + 1 s.d. Note that at this point “cell” can refer to single cells or cell conglomerates.

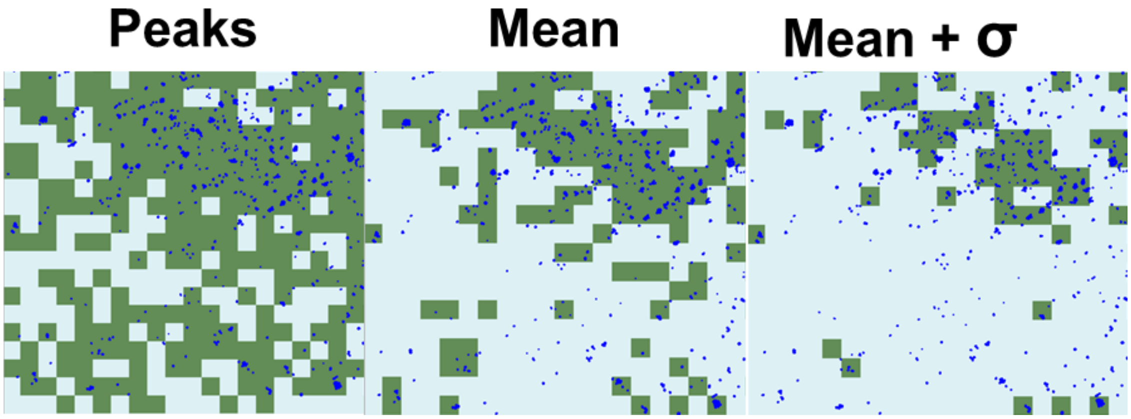

Figure S8. PreScan Outcomes at Various Thresholds versus Hoechst-Stained Cells as Ground Truth. Comparison of different PreScan thresholds, using *m/z* 281.25 (FA18:1), and outcomes at a 200-µm pixel size (green) with Hoechst-stained cell nuclei (blue). This visual example complements the previous figure. Hoechst stain was performed after MSI and co-registered. "Peaks" implies that each *m/z* feature was selected to create the mask that was used for the subsequent DeepScan. "Mean" indicates the peak threshold established at the average intensity of *m/z* = 281.25 in all spectra, and "mean + σ", the most restrictive setting, refers to the threshold being adjusted to the mean intensity plus one standard deviation (σ) of all pixels in this measurement.

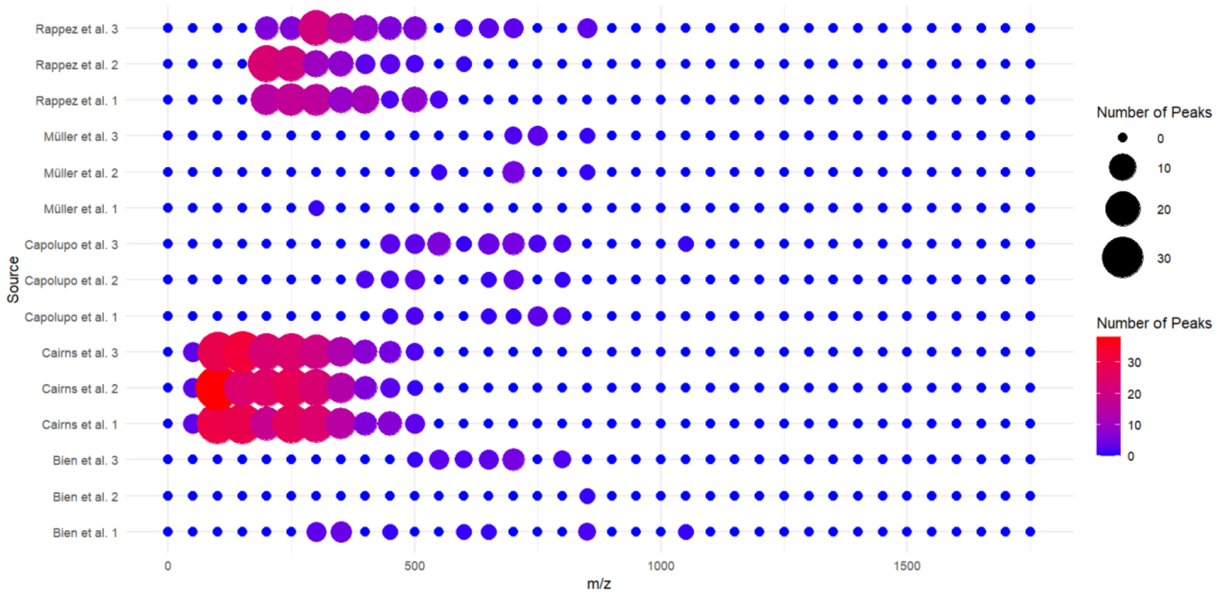

Figure S9. Mass Ranges in State-of-the-Art Single Cell MSI Technologies. For all publicly available single cell .imzMLs via Metaspace we picked n=3 per research group and compared the number of peaks that correlated with HMDB endogenous *m/z* intervals found in 100 randomly sampled pixels. This random sampling was done to account for different dataset sizes. Note that we are comparing different spatial resolutions here, which will impact the number of hits, as sensitivity decreases with small pixel size. Bien et al. and Müller et al. used <5 µm spatial resolution. Also likely differences between measurement devices are not accounted for here. Aim of this plot was to show that PRISM-MS (Cairns et al.) covers true small molecules with *m/z* < 200 using MALDI imaging and does not focus on lipids. Each data point represents an *m/z* interval of 50.

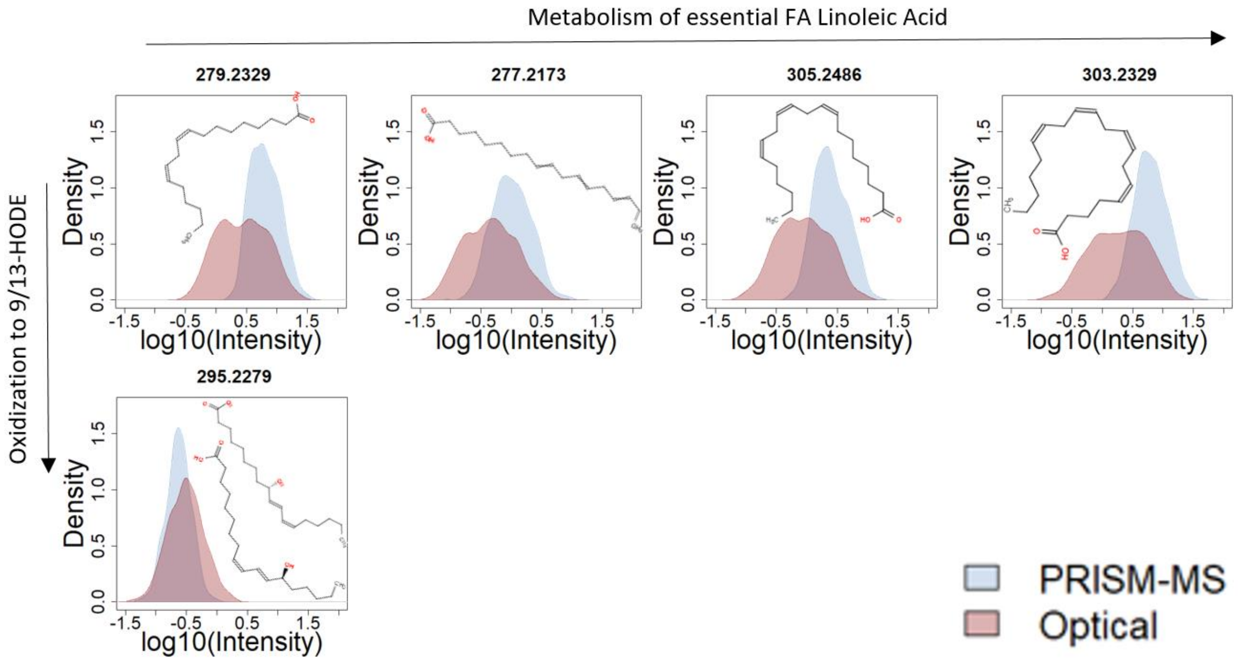

Figure S10. Comparative Analysis of Linoleic Acid Metabolism and Oxidation in "Optically-Guided" and PRISM-MS Workflows. An emulated optically-guided workflow (red), where the slide was placed in a sterile hood for 30 min at RT prior to measurement, led to a decrease in linoleic to arachidonic acid conversion pathway metabolites[1] from the essential unsaturated fatty acid linoleic acid (*m/z* = 279.2329 $[M-H]^-$) to arachidonic acid (*m/z* = 303.2329 $[M-H]^-$) via gamma-linoleic acid (*m/z* = 277.2173 $[M-H]^-$) and di-homo-gamma-linoleic acid (*m/z* = 305.2486 $[M-H]^-$). An increase in oxidized metabolites 9- and 13-hydroxy-octadecadienoic acid (9/13 HODE, *m/z* = 295.2279 $[M-H]^-$) was observed in the optically guided workflow (red) compared to the PRISM-MS workflow (blue). PRISM-MS avoids a high-resolution optical image before starting the targeted MALDI MSI run. The analysis uses a density function, represented as a smoothed histogram through kernel density estimation on the y-axis, plotted against $\log_{10}$ of peak intensities. Data from n=3 slides was combined. Consistently, signals for linoleic acid and its metabolites were preserved using PRISM-MS, whilst the amounts of oxidation products was reduced.

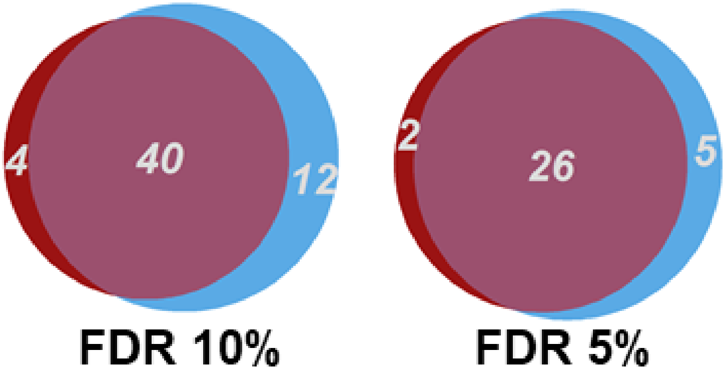

Figure S11. Metaspace Annotations for PRISM-MS and Optical Guidance. Venn diagrams comparing the number of metabolite annotations in METASPACE at 10% and 5% FDR, for PRISM-MS (blue) and optically-guided MALDI MSI (red).

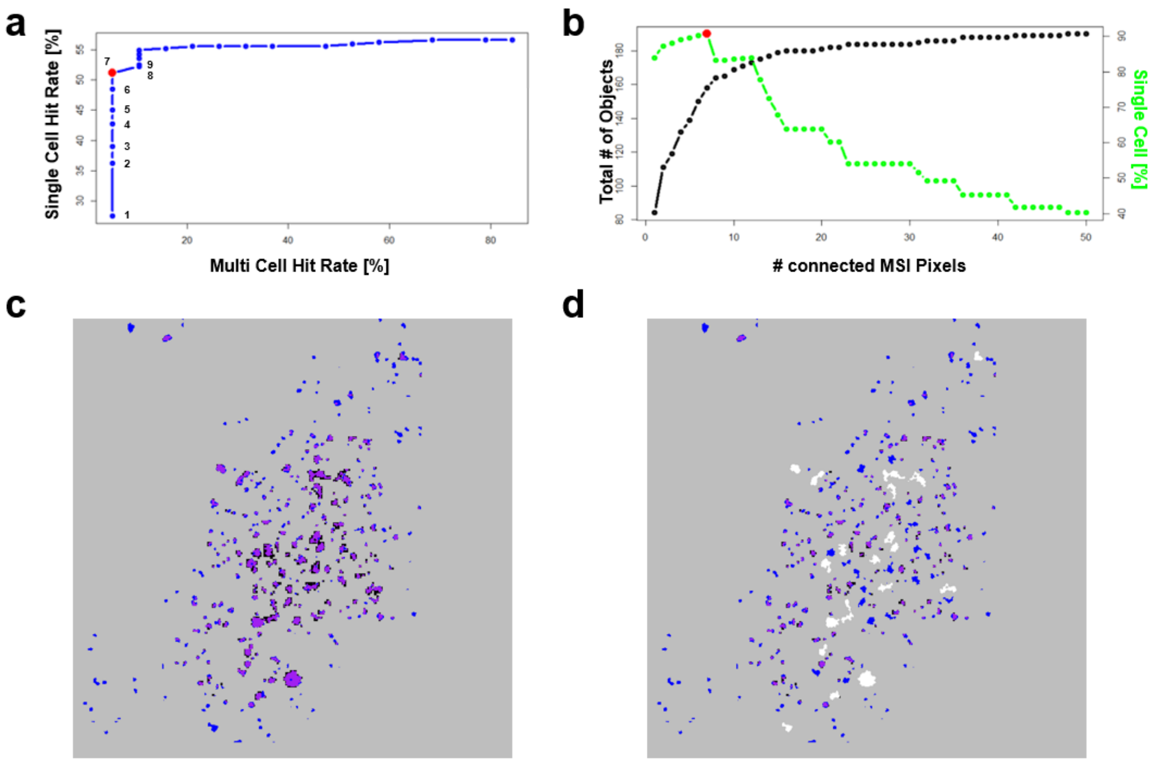

Figure S12. Comparative Analysis of Single- and Multi-Cell Capture Rates Using MSI Processing Pipeline with Optimal Pixel Connectivity. Evaluation of data processing pipeline following the DeepScan (20 µm), which i) clusters pixels into "cells"/objects, ii) distinguishes single cells (single cell hit rate (SC-HR)) from cell aggregates/conglomerates (multi cell hit rate (MC-HR)) and iii) removes aggregates from consideration. A single SIMA9 cell is defined as having a coherent Hoechst-stained area ≤100 connected image pixels (~400 µm²)[2] in a fluorescence image with 2 µm pixel size. Nuclei comprise >80% of the SIMA9 cell area that has a ~20 µm diameter[3]. PreScan data (200 µm) was acquired in negative ion mode MSI from a low seeding-density SIMA9 cell culture using intensity threshold mean + 1 s.d. and a spot dilation factor 1. The feature for cell detection was FA18:1 (*m/z* = 281.25). **a)** SC-HR, *i.e.,* the percentage of Hoechst staining-defined single cells captured by the PRISM-MS processing pipeline, was compared with the MC-HR, *i.e.,* the percentage of cell aggregates (anything larger than single cell) captured. Data points in the trajectory correspond to [N] connected MSI pixels that constitute one single cell. Here, the optimum was n = 7 connected MSI pixels (red point), for which single cell identification was maximized whilst capture of cell aggregates was minimized. Cell aggregates were falsely classified as single cells, *i.e.*, MC-HR started to increase, when 8 or more connected pixels were assumed as single cells. **b)** The number N of connected MSI pixels is plotted against the total number of objects captured (single cell plus cell aggregates; black) and the percentage of single cells within all objects (single cell plus cell aggregates; green) with the optimal point, n=7, indicated in red. At this point 90.7 % of all detected cells were true single cells. **c)** Overlay image showing Hoechst staining (blue), MSI signals (black) and overlapping areas (purple) without setting a connected pixel threshold. **d)** Same coloring scheme but applying the maximum of n=7 connected pixels to label multicellular objects (white). These larger objects are excluded from the data to enable single cell analysis.

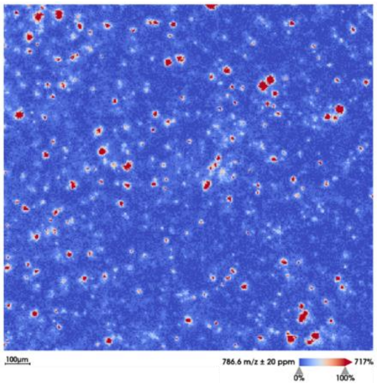

Figure S13. MALDI MS Imaging of DOPC-GUVs. MALDI MS Imaging of DOPC (*m/z* = 786.6) at 5 µm lateral step-size.

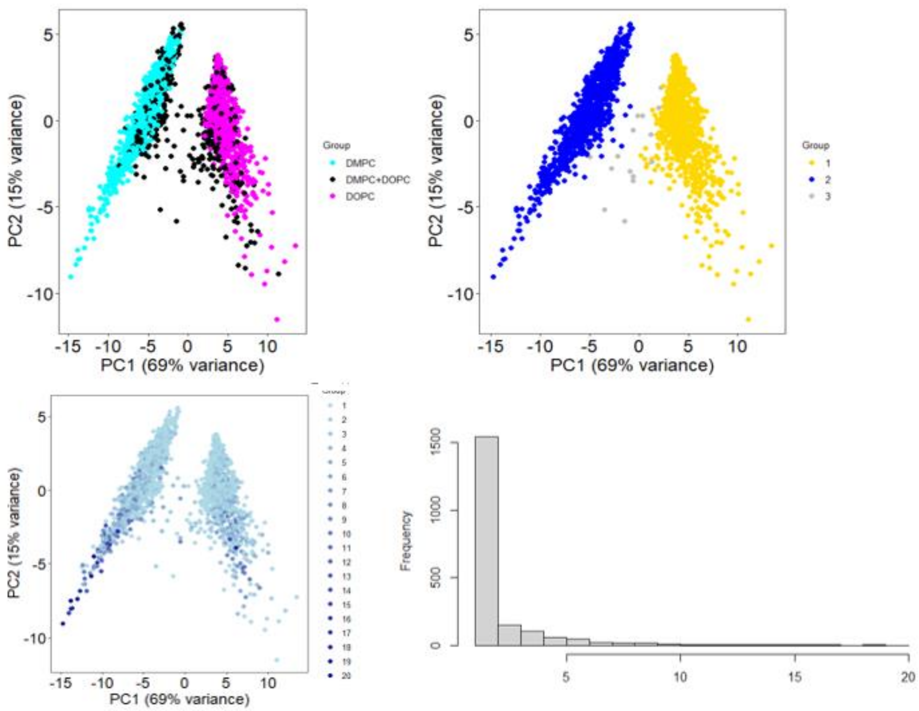

Figure S14. Analysis of GUV Sizes and M3C Method Top Left: Principal Component Analysis (PCA) of Giant Unilamellar Vesicles (GUVs), imaged at 20 µm pixel size and data processed by clustering pixels into vesicles with labels based on GUV composition: DOPC only, DMPC only or mixed DMPC+DOPC. Top Right: Monte Carlo Consensus Clustering (M3C) analysis classifies GUVs into three groups, *i.e.,* being positive for DMPC (1), DOPC (2) or both (3). Vesicles classified as positive for both (3) were mostly two vesicles in close proximity that were not separated at 20 µm lateral step size (compare zoom-in **Figure 2f**). Bottom Left: PCA illustrating GUV sizes, displayed as numbers of connected pixels for each vesicle, with darker shades of blue representing larger vesicles up to 20 connected pixels. This analysis suggests that vesicle size has no impact on cluster assignment in our defined system. Bottom Right: Histogram indicating variability in GUV sizes, with the majority of them being displayed as single 20 µm pixels.

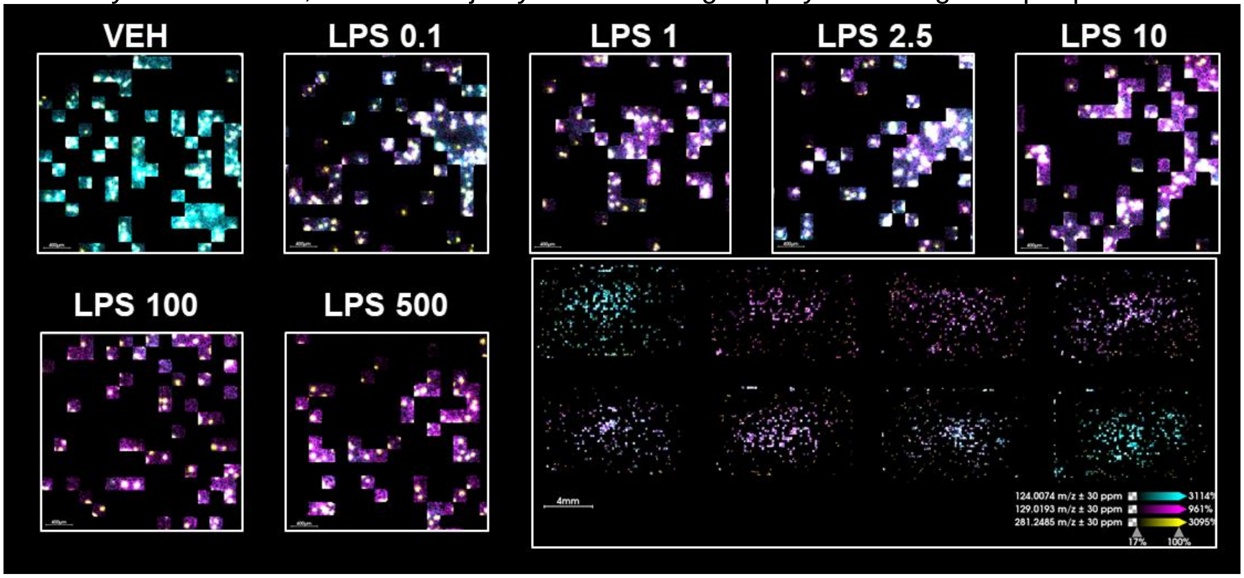

Figure S15. PRISM-MS DeepScan of LPS-treated SIMA9 Cells: Spatial Analysis of Itaconate, Taurine, and FA 18:1. MALDI Imaging DeepScan of SIMA9 cells treated with 0 to 500 ng/mL lipopolysaccharide (LPS) at 20 µm pixel size following a PreScan at 200 µm lateral step-size using a spot dilation of 1 and thresholding of mean + s.d. [σ]. Lower right panel: overview of the entire slide, where concentration-dependence of itaconate- (magenta) and taurine-levels (cyan) is visible. Displayed are itaconate (*m/z* 129.0193; magenta), taurine (*m/z* 124.007; cyan), and the fatty acid FA 18:1 (*m/z* 281.2485; yellow).

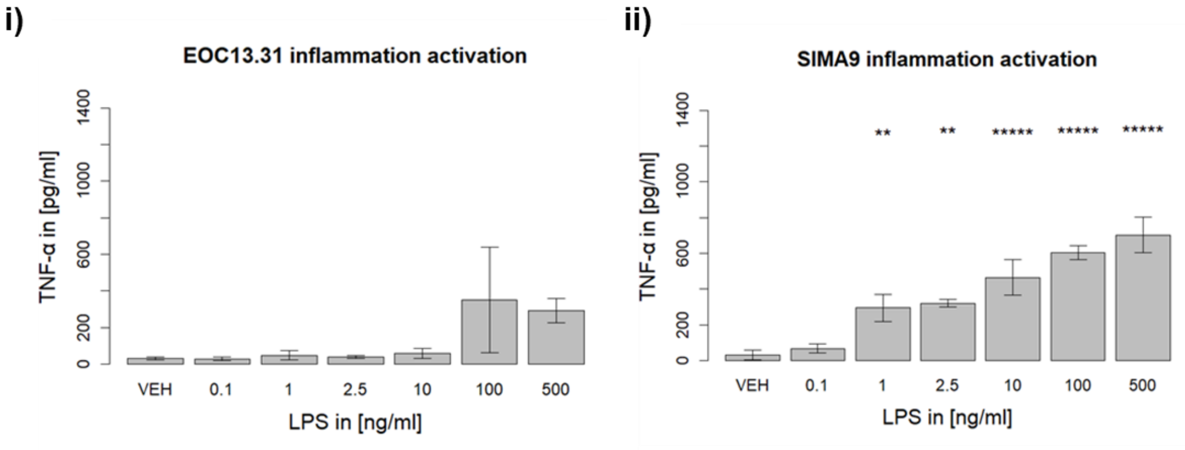

Figure S16. TNF-α Levels in EOC13.31 and SIMA9 Cells Post-LPS Stimulation. TNF-α levels secreted by i) EOC13.31 and ii) SIMA9 cells were measured via sandwich ELISA after treatment with vehicle (VEH; 0 ng/mL) or with varying LPS concentrations (0.1, 1, 2.5, 10, 100,

500 ng/mL) for 20h. Data represent the mean ± s.d. One-way ANOVA **** $p < 0.0001$, ** $p < 0.01$, * $p < 0.05$ to compared to VEH. N=3 per condition.

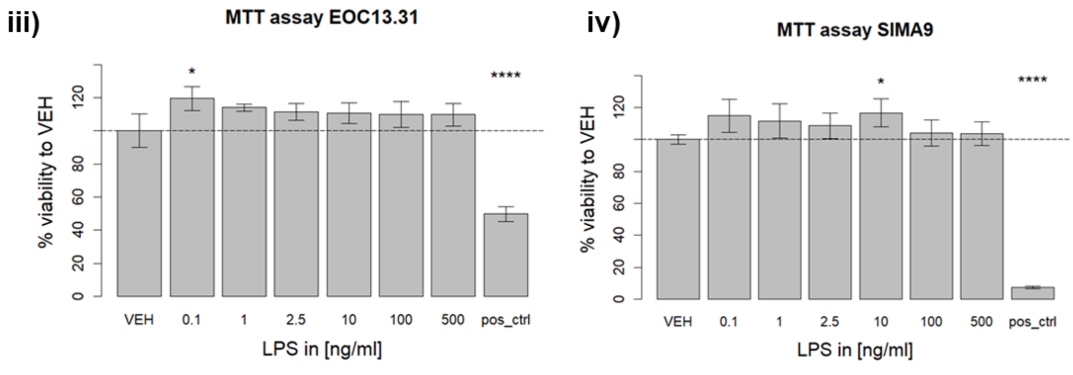

Figure S17. MTT Assay for EOC13.31 and SIMA9 Cells Post-LPS Stimulation. MTT assay of i) EOC13.31 and ii) SIMA9 cells. MTT metabolic activity was measured after treatment with vehicle (VEH; 0 ng/mL) or with varying LPS concentrations (0.1, 1, 2.5, 10, 100, 500 ng/mL) for 20h. 1% Triton X-100 was used as positive control for reduced cell viability (pos_ctrl). Data represent the mean ± s.d. One-way ANOVA **** $P < 0.0001$, ** $P < 0.01$, * $P < 0.05$ compared to VEH. N=3 per condition.

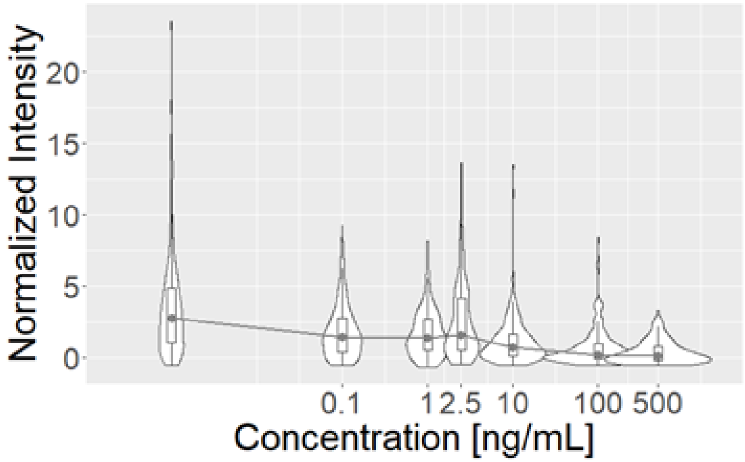

Figure S18. Taurine Levels after LPS Treatment in SIMA9 cells. Violin plot of taurine (*m/z* = 124.01) ion intensities after data processing to assign pixel clusters to single cells. Mean ion intensities for taurine, normalized to the internal standard Trp-D5 and Z-score-standardized per cell, correlated with LPS concentrations (0 ng/mL to 500 ng/mL). Same data as shown in **Figure S 15** and corresponding to the violin plot for itaconate in **Figure 3b**.

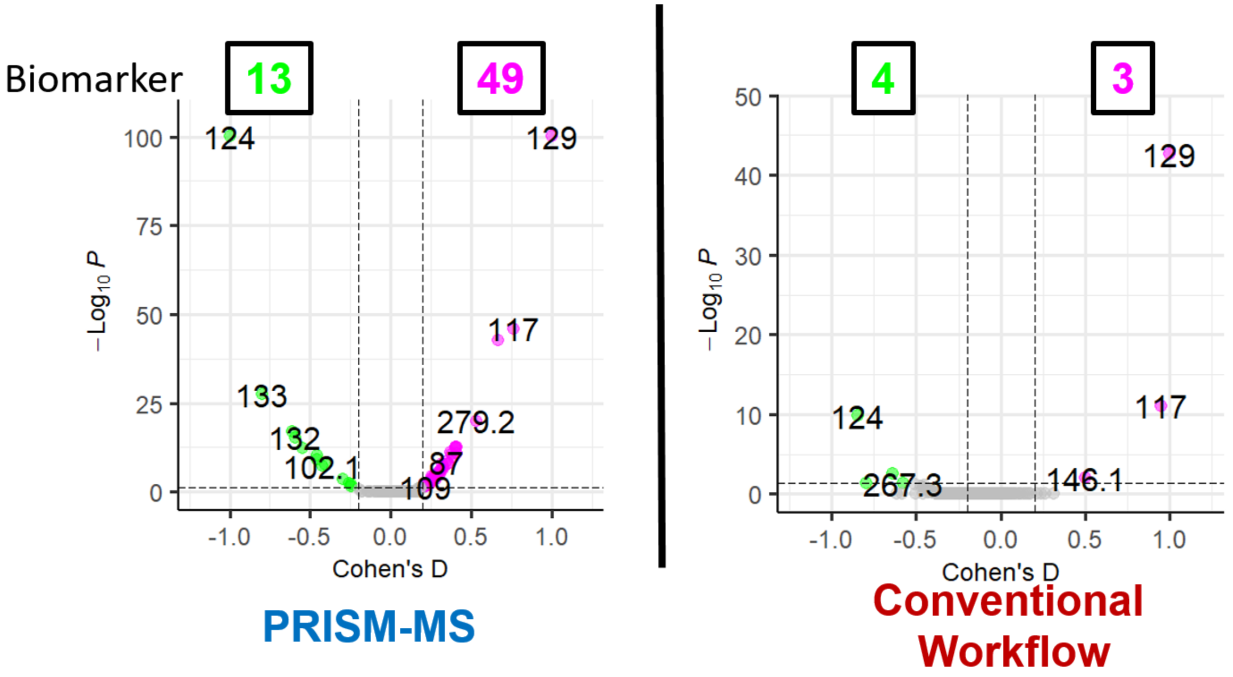

Figure S19. PRISM-MS with M3C enables Identification of more Candidate Activation Biomarkers. SIMA9 cells were treated with LPS. Left: PRISM-MS with M3C resulted in 13 and 49 candidates for resting- and activation-specific biomarkers, respectively. Right: The same computational pipeline but for a conventional workflow, in which acquisition of a microscopic image was emulated, resulted in 4 and 3 candidates for resting- and activation-specific biomarkers, respectively. Two possible reasons: First, all ion intensities and, thus, detectability of metabolites is lower using the conventional workflow, and finding differences becomes harder in general. Second, the biomarkers used to define active SIMA9 cells (itaconate) or resting SIMA9 cells (taurine) are also lowered, which consequently makes finding and defining these subpopulations more challenging.

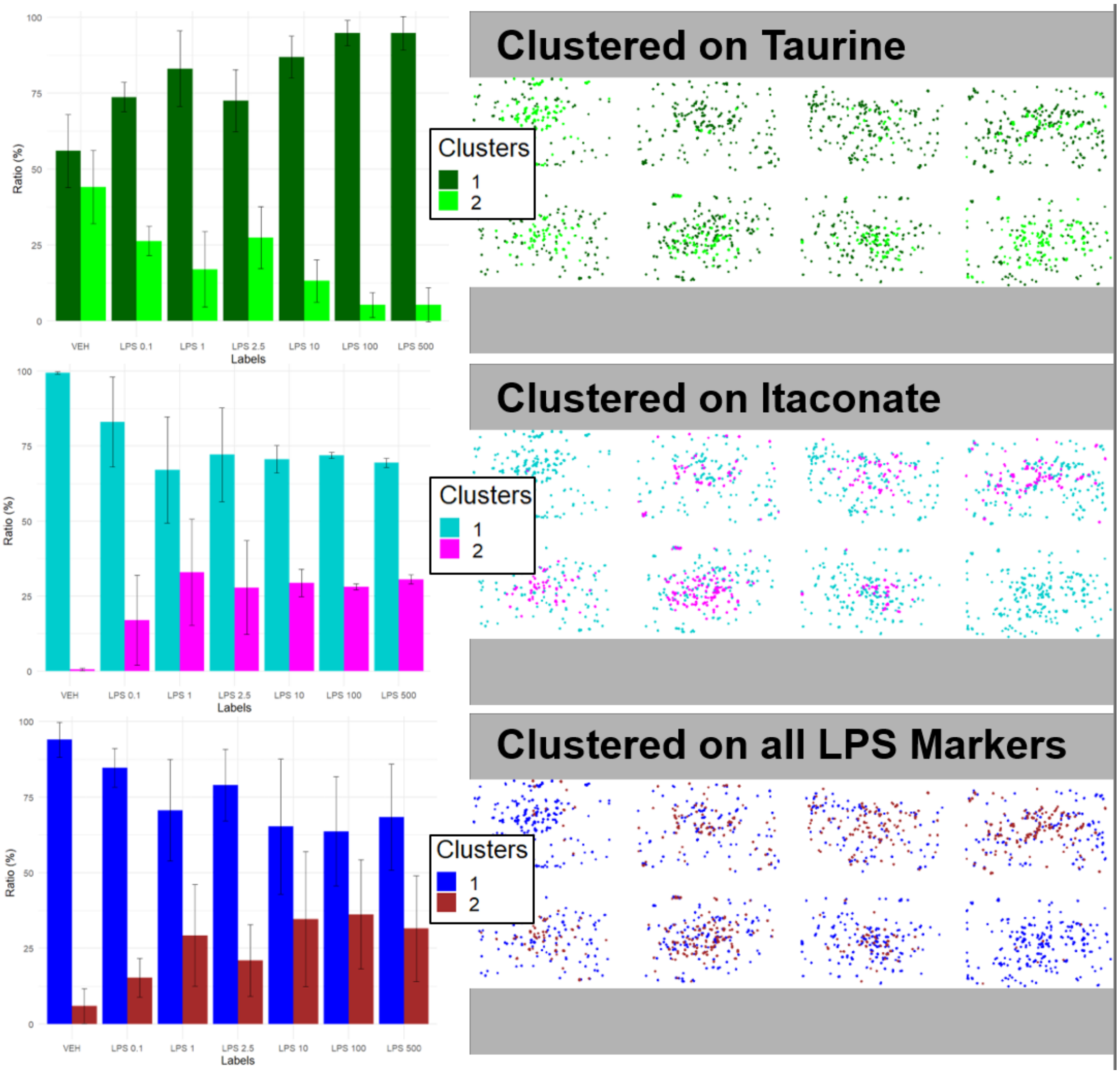

Figure S20. M3C Analysis of Taurine (top), Itaconate (middle), and Combined LPS-activation Markers (bottom) in SIMA9 Cells. SIMA9 cells were stimulated with LPS and candidate markers for LPS activation were identified as in Figure 3a. Note that population concentration-responses were similar for M3C with itaconate *m/z* versus M3C based on all LPS activation-specific features. M3C with taurine *m/z* led to an inverse population response compared to itaconate. These M3Cs were used for label creation taur+ita+, taur-ita+, taur+ita-, and taur-ita-. Subsequently, we improved data quality by removing taur-ita- cells from consideration. The nature of these cells is unclear: they may have died, burst open have released metabolites, may be non-reactive to the stimulus in general or be in some other stage of the cell cycle whilst still being detected using the cell marker FA18:1 (*m/z* = 281.25). Left: N=3 separate slides were analyzed to produce the boxplots (percentage of cells for each treatment sorted into specific clusters). Right: Corresponding M3C clustered PRISM-MS DeepScans for an exampe slides.

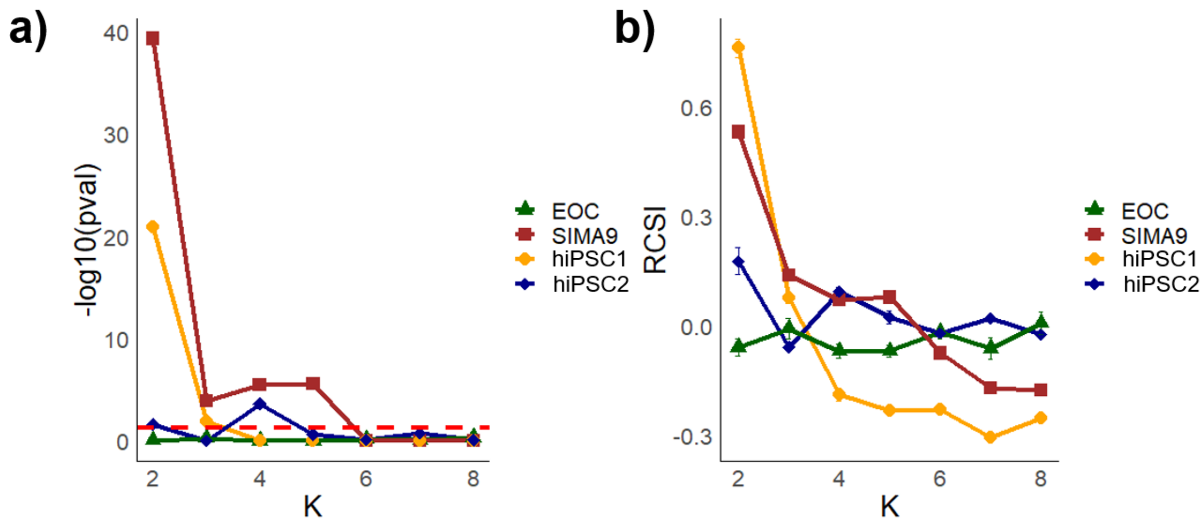

Figure S21. P-Value Analysis and Relative Cluster Stability Index (RCSI) across Microglia-like Cell Lines. Left panel: M3C clustering on itaconate (*m/z* = 129.02) and p-value analysis for cell lines SIMA9 and EOC as well as the hiPSC lines hiPSC1 and hiPSC2cells for different cluster numbers k. For each cell line and for each number of clusters (k) the test was repeated 200 times to determine if the population of cells was significantly different from a uniform/homogeneous distribution without subpopulations. For EOC cells, no k exceeded the significance threshold of 0.01, indicating that EOC cells did not have active subpopulations and were not responsive to LPS. This was expected, since EOC cells lack toll-like receptor 4 (TLR4). SIMA9, hiPSC1 and hiPSC2 cells showed significant subpopulations, for multiple k. Right panel: Since the p-value cannot be compared directly with another p-value to determine the optimal k, a second metric, the Relative Cluster Stability Index (RCSI), was used to determine this. RCSI measures the stability of clusters to find an optimal number of clusters (k). RCSI analysis across different cluster numbers for SIMA9, hiPSC1, and hiPSC2cells suggested k=2 as the optimal cluster number based on stability and significance according to p-values. For hiPSC2, both k=2 and k=4 were candidates based on p-value significance; however, k=2 was chosen due to higher RCSI, indicating a stronger cluster stability. For SIMA9 and hiPSC1 cells k=2 was significant and had the highest cluster stability.

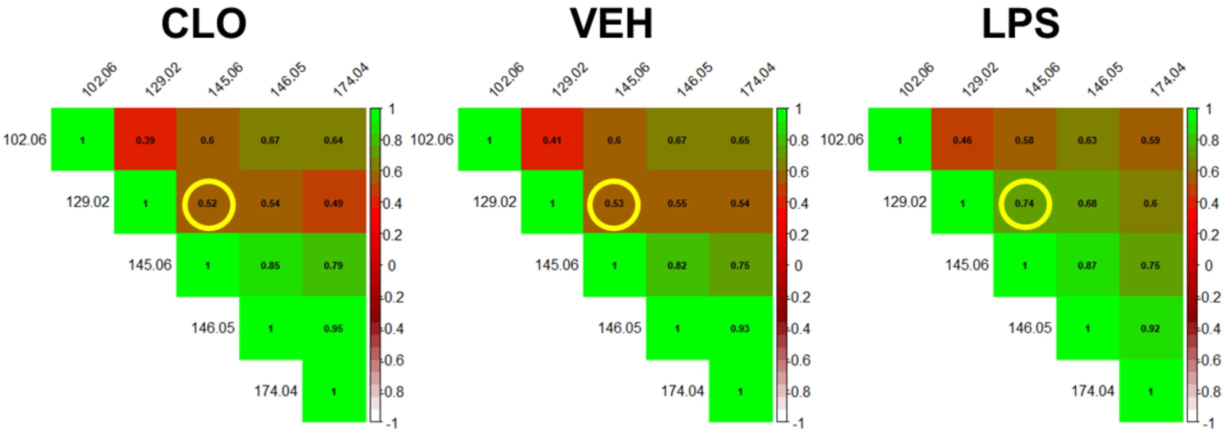

Figure S22. Correlation Change of Itaconate and Glutamine across Treatment Conditions for Hippocampal Slice Cultures. Pearson intensity correlation plots between *m/z* values for different treatment conditions (PBS/vehicle (VEH), clodronate (CLO), LPS) of rat hippocampal slice cultures. The measurement was performed 5 times on different days. Each experiment had all three conditions (VEH, CLO & LPS) in triplicates, resulting in a total of 45 sections that were analysed. The total amount of pixels for these experiments were 1.2million. The yellow circle marks the correlation between itaconate (*m/z* = 129.02) and glutamine (*m/z* = 145.06) across the different conditions. While weak correlations were observed in CLO (0.52) and VEH (0.53) treated tissue, a strong correlation was noted in LPS-treated tissue (0.74), indicating colocalization in this condition. This implied that higher itaconate intensity was associated with higher glutamine intensity in LPS-treated tissue, unlike in VEH and CLO treatments. Other features shown include GABA (*m/z* = 102.06), glutamate (*m/z* = 146.05), and N-acetyl-aspartate (*m/z* = 174.04).

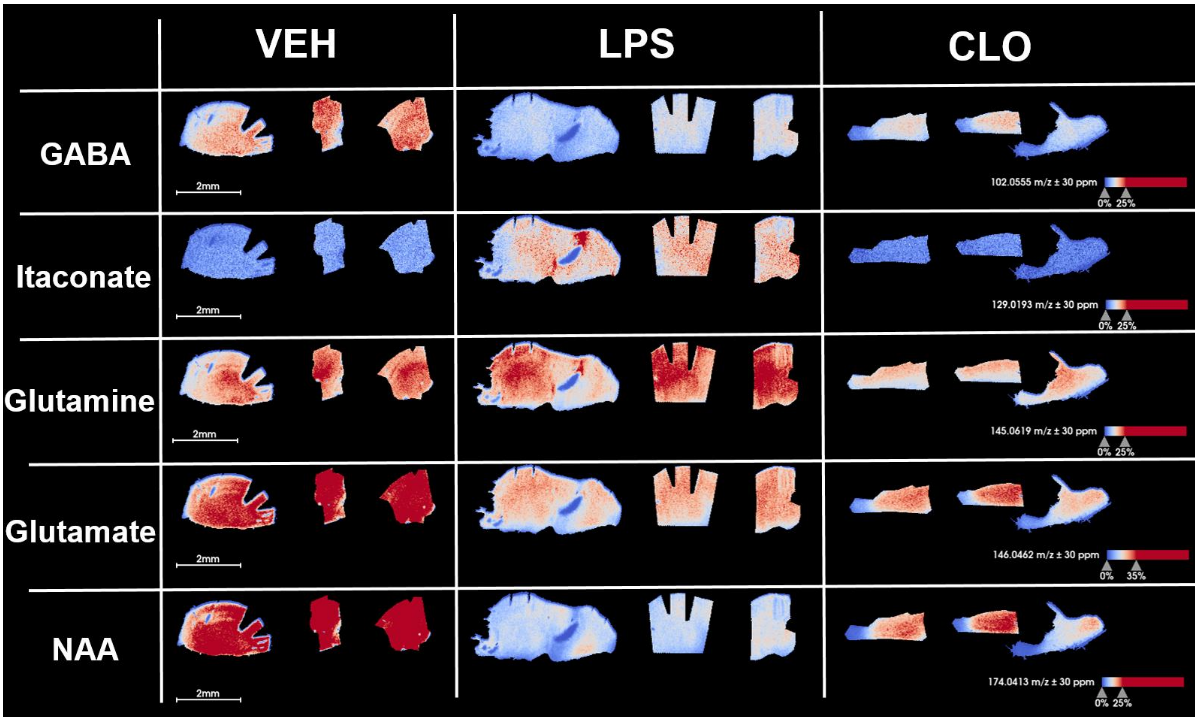

Figure S23. MALDI MSI Analysis of Gamma-Amino-Butyric Acid (GABA; *m/z* 102.055), Itaconate (*m/z* 129.019), Glutamine (*m/z* 145.062), Glutamate (*m/z* 146.046), and N-Acetyl-Aspartate (NAA; *m/z* 174.041). MALDI MS Imaging in negative ion mode of rat hippocampal slice cultures (whole slices or fragments thereof), treated with PBS (vehicle; VEH) or 1 µg/mL LPS or 100 µg/mL clodronate (CLO), at 20 µm pixel size. Ion images show metabolites found to be significantly different (as determined by volcano plot) between VEH and LPS in Slice Cultures as well as in cell cultures (see **Supplementary Table 1**).

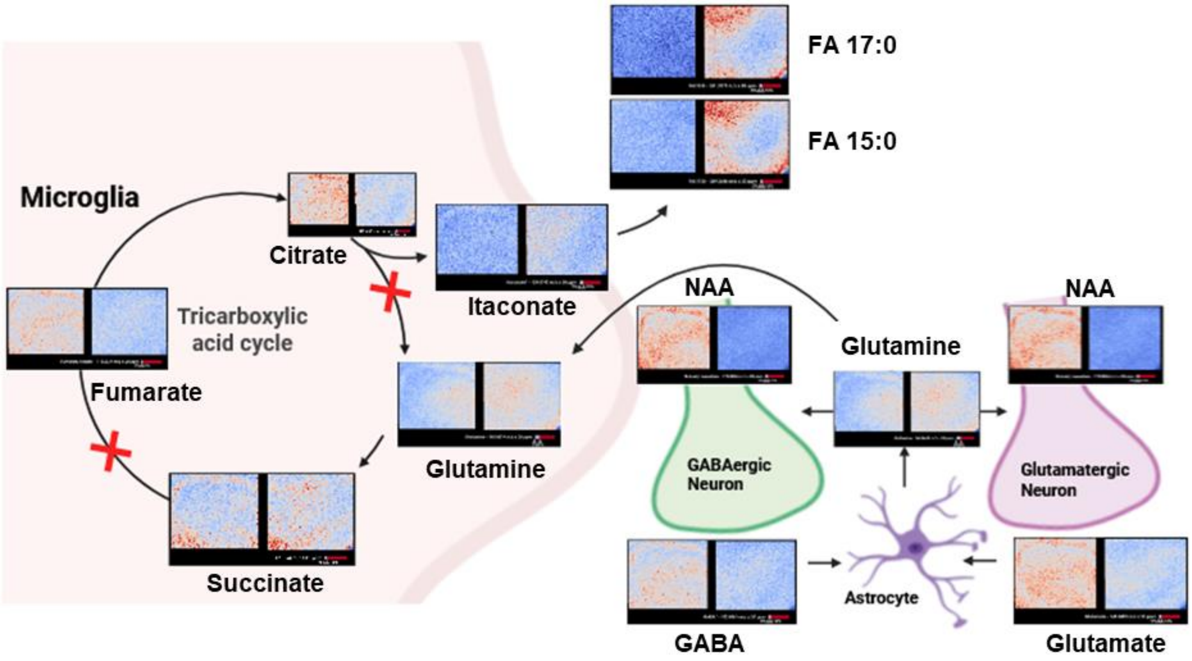

Figure S24. Hypothetical of microglia tapping into metabolite shuttling between astrocytes, and excitatory/inhibitory neurons size. MALDI MS Imaging of rat hippocampal slice cultures, treated with 0 ng/mL (vehicle; VEH) or 500 ng/mL (LPS), at 5 µm pixel size. Ion images of TCA cycle metabolites and the hypothesized pathway pulling glutamine from neighboring cells while also explaining N-acetyl aspartate (NAA) depletion. In each image, the left panel is VEH control and the right panel is LPS-treated tissue. The ion images represent: Citrate $[M-H]^-$ (*m/z* 191.02), Itaconate $[M-H]^-$ (*m/z* 129.02), Glutamine $[M-H]^-$ (*m/z* 145.06), Succinate $[M-H]^-$ (*m/z* 117.02), Fumarate $[M-H]^-$ (*m/z* 115.00), FA15:0 $[M-H]^-$ (*m/z* 241.22), FA17:0 $[M-H]^-$ (*m/z* 269.25), GABA* $[M-H]^-$ (*m/z* 102.06), glutamate $[M-H]^-$ (*m/z* 146.05) and NAA $[M-H]^-$ (*m/z* 174.04).

### Supplementary Tables

Table S1. Overview of Significant Features, as Determined via Volcano Plot, for Rat Hippocampal Slice Cultures. Names marked with * have alternative annotations or isoforms that cannot be distinguished via MS/MS or high resolution mass spectrometry. Annotations were done on timsTOFflex imaging data via METASPACE using the HMDB and KEGG data bases. Afterwards the accurate masses were determined using an FTICR for validation. All ions detected as $[M\text{-}H]^-$. The complete table of METASPACE annotations is an additional supplementary data .csv file.

| Name | FDR KEGG | *FDR HMDB* | Sum formula | *m/z* theo. | *m/z* FTICR | ppm FTICR |
|---|---|---|---|---|---|---|
| GABA* | 5 | 5 | $[C_4H_9NO_2 - H]^-$ | 102.0561 | 102.05609 | 0.10 |
| Fumarate/Malate | 5 | 5 | $[C_4H_4O_4 - H]^-$ | 115.0037 | - | - |
| Taurine | 5 | 5 | $[C_2H_7NO_3S - H]^-$ | 124.0074 | 124.00736 | 0.32 |
| Itaconate* | 5 | 5 | $[C_5H_6O_4 - H]^-$ | 129.0193 | 129.01934 | 0.31 |
| N-Acetyl-Alanine | 10 | 10 | $[C_5H_9NO_3 - H]^-$ | 130.051 | 130.05093 | 0.54 |
| Ornithine | 10 | 10 | $[C_5H_{12}N_2O_2 - H]^-$ | 131.0826 | 131.08258 | 0.15 |
| Adenine | 5 | 5 | $[C_5H_5N_5 - H]^-$ | 134.0472 | 134.04717 | 0.22 |
| Hypoxanthine | 5 | 5 | $[C_5H_4N_4O - H]^-$ | 135.0312 | 135.0312 | 0.00 |
| Glutamine | 5 | 5 | $[C_5H_{10}N_2O_3 - H]^-$ | 145.0619 | 145.06182 | 0.55 |
| Glutamate | 5 | 5 | $[C_5H_9NO_4 - H]^-$ | 146.0459 | 146.04584 | 0.41 |
| Glycerol-Phosphate | 5 | 5 | $[C_3H_9O_6P - H]^-$ | 171.0064 | 171.00643 | 0.18 |
| N-Acetyl-Aspartate | 5 | 5 | $[C_6H_9NO_5 - H]^-$ | 174.0408 | 174.04079 | 0.06 |
| Kynurenate | 5 | 5 | $[C_{10}H_7NO_3 - H]^-$ | 188.0353 | 188.03534 | 0.21 |
| Glycerol-3-Phospho-ethanolamine | 5 | 5 | $[C_5H_{14}NO_6P - H]^-$ | 214.0486 | 214.04864 | 0.19 |
| FA(15:0) | 5 | 5 | $[C_{15}H_{30}O_2 - H]^-$ | 241.2173 | 241.21735 | 0.21 |
| Cytidine | 10 | 10 | $[C_9H_{13}N_3O_5 - H]^-$ | 242.0782 | - | - |
| FA(17:0) | 5 | 5 | $[C_{17}H_{34}O_2 - H]^-$ | 269.2486 | 269.24857 | 0.11 |
| FA(18:1) | 5 | 5 | $[C_{18}H_{34}O_2 - H]^-$ | 281.2486 | 281.24863 | 0.11 |
| Arachidonate | 5 | 5 | $[C_{20}H_{32}O_2 - H]^-$ | 303.2329 | 303.23283 | 0.23 |
| Glutathione | 5 | 5 | $[C_{10}H_{17}N_3O_6S - H]^-$ | 306.0765 | 306.07636 | 0.46 |
| FA(22:5) | 5 | 5 | $[C_{22}H_{34}O_2 - H]^-$ | 329.2486 | 329.24846 | 0.43 |
| AMP | 5 | 5 | $[C_{10}H_{14}N_5O_7P - H]^-$ | 346.0558 | 346.05567 | 0.38 |
| IMP | 5 | 5 | $[C_{10}H_{13}N_4O_8P - H]^-$ | 347.0398 | 347.03979 | 0.03 |

**Table S2. Summary of MS2-based Formula Identification for Metabolites in Hippocampal Slice Cultures via SIRIUS.** Overview of MS2 fragmentation analysis results for metabolite hits annotated via SIRIUS[4] performed on rat hippocampal slice cultures. For each precursor, the sum formula with the highest Sirius Score is displayed. The same samples were also analyzed by FT-ICR MSI for accurate mass determination. The sum formulas marked in red got the highest SIRIUS scores for the precursor, whilst showing a different sum formula than the METASPACE database annotations and/or FTICR exact mass measurement. All precursors marked with * are listed in **Table S3**. These precursors additionally got compound annotations via Database Search and received CSI:FingerID Scores for this.

| ***m/z* Precursor** | **Sirius Score** | ***Number of Peaks*** | **Explained Peaks** | **Sum formula** | **Median mass error fragments (ppm)** |
|---|---|---|---|---|---|
| 102.06 | 0.99 | 3 | 2 | C4H9NO2 | -0.2 |
| 115.00 | 6.88 | 3 | 2 | C4H4O4 | -8.4 |
| 124.01* | 10.72 | 10 | 4 | C2H7NO3S | -3.7 |
| 129.02 | 17.28 | 6 | 2 | C5H6O4 | -4.7 |
| 130.05* | 13.81 | 15 | 5 | C5H9NO3 | 0.0 |
| 134.05* | 14.73 | 11 | 6 | C5H5N5 | 0.6 |
| 135.03* | 15.42 | 13 | 5 | C5H4N4O | -0.9 |
| 145.06* | 38.94 | 33 | 8 | C5H10N2O3 | -0.7 |
| 146.05* | 27.21 | 39 | 7 | C5H9NO4 | 0.3 |
| 174.04* | 51.60 | 26 | 10 | C6H9NO5 | -2.7 |
| 241.22 | 3.49 | 5 | 1 | C15H30O2 | -0.1 |
| 242.08* | 30.34 | 36 | 12 | C7H11N6O4 | 3.8 |
| 281.25 | 3.46 | 3 | 1 | C18H34O2 | -0.5 |
| 303.23* | 13.69 | 24 | 5 | C20H32O2 | 1.8 |
| 306.08* | 21.68 | 33 | 20 | C10H17N3O6S | -3.1 |
| 329.25* | 6.65 | 21 | 3 | C22H34O2 | 1.3 |
| 346.06* | 10.24 | 15 | 4 | C9H18NO11P | -2.5 |
| 347.07* | 8.24 | 14 | 4 | C9H17O12P | -3.3 |

#### Table S3. Summary of MS2-based Metabolite Identification on Slice Culture via SIRIUS.

Metabolites were confirmed via MS2 fragmentation analysis and KEGG database search. Metabolites marked in green are the correct sum formulas (as opposed to the falsely annotated ones in **Table S2** using just SIRIUS scoring) that match with MS2 database search as well as imaging METASPACE annotations and exact mass measurements via FTICR. The reason for this mismatch is that they had the second highest SIRIUS scores[4] for each precursor in **Table S2**. Full data is available in the attached Sirius_Supp.xlsx file.

| ***m/z* Precursor** | **Confidence Score** | ***CSI:FingerID*** | **Predicted Fingerprints** | **Name** | **Sum formula** |
|---|---|---|---|---|---|
| 124.01 | 0.415 | -3.29 | 1 | Taurine | C2H7NO3S |
| 130.05 | 0.435 | -12.77 | 2 | N-Acetyl-L-Alanine | C5H9NO3 |
| 134.05 | 0.337 | -10.57 | 1 | Adenine | C5H5N5 |
| 135.03 | 0.257 | -9.02 | 1 | Hypoxanthine | C5H4N4O |
| 145.06 | 0.167 | -27.71 | 2 | Glutamine | C5H10N2O3 |
| 146.05 | 0.271 | -12.24 | 1 | Glutamate | C5H9NO4 |
| 174.04 | 0.573 | -4.64 | 2 | N-Acetyl-L-aspartic acid | C6H9NO5 |
| 242.08 | 0.104 | -164.42 | 3 | Cytidine | C9H13N3O5 |
| 303.23 | 0.079 | -17.65 | 1 | Arachidonate | C20H32O2 |
| 306.08 | 0.136 | -12.34 | 2 | Glutathione | C10H17N3O6S |
| 329.25 | 0.063 | -64.25 | 1 | Docosapentaenoic acid | C22H34O2 |
| 346.06 | 0.354 | -8.76 | 4 | AMP | C10H14N5O7P |
| 347.04 | 0.255 | -12.19 | 7 | IMP | C10H13N4O8P |

### Supplementary Datasets provided as separate folders

Dataset S1. Sirius file of MSMS analysis of peaks from Supplementary Table 1, 2 & 3

### Supplementary Datasets provided as separate Excel Tables

Dataset S2. Annotations via Metaspace for KEGG & HMDB

Dataset S3. Summary of Metabolic Changes after LPS Stimulation for SIMA9, EOC, hiPSC1, hiPSC2 and all Six Slice Cultures
